# Pan-transcriptome reveals a large accessory genome contribution to gene expression variation in yeast

**DOI:** 10.1101/2023.05.17.541122

**Authors:** E. Caudal, Victor Loegler, F. Dutreux, N. Vakirlis, E. Teyssonnière, C. Caradec, A. Friedrich, J. Hou, J. Schacherer

## Abstract

Gene expression is an essential step in the translation of genotypes into phenotypes. However, little is known about the transcriptome architecture and the underlying genetic effects at a species-level. Here, we generated and analyzed the pan-transcriptome of ∼1,000 yeast natural isolates across 4,977 core and 1,468 accessory genes. We found that the accessory genome is an underappreciated driver of the transcriptome divergence. Global gene expression patterns combined with population structure show that the heritable expression variation mainly lies within subpopulation-specific signatures, for which the accessory genes are overrepresented. Genome-wide association analyses consistently highlight that the accessory genes are associated with proportionally more variants with larger effect sizes, illustrating the critical role of the accessory genome on the transcriptional landscape within and between populations.

## Introduction

Gene expression is the primary step of a process by which information encoded in the genome is converted into traits. Genetic variants affecting gene expression levels contribute to the phenotypic diversity^1–4^. The dissection of the genetic regulation of the different molecular intermediates leading to the final phenotypes is therefore essential for understanding many aspects of biology. Genetic variants or loci associated with gene expression variation (*i.e.*, expression Quantitative Trait Loci, eQTL) have been identified via linkage and genome-wide association mapping in various organisms, uncovering general mechanisms of transcription regulation^5–16^. Genetic variants underlying gene expression variation can be located close or further to the affected gene and are hence considered as *cis*-acting (local eQTL) or *trans*-acting (distant eQTL), respectively. All these studies highlighted the fact that local eQTL have larger effects on gene expression than distant eQTL^5–7^. However, a gene is often regulated by multiple distant eQTL and this set of *trans*-regulatory variants tend to explain a larger fraction of the variance overall than its *cis*-eQTL^6^.

In recent years, large-scale transcriptomic surveys have been initiated in multiple model and non-model systems^5, 7, 12, 14, 17^, most notably the GTEx project in humans which covers gene expression data across 49 tissues from up to 838 individuals^12^. These studies were extremely insightful into the tissue-specific gene expression regulations and extensively catalogued *cis*-acting variants acting across tissues^12^. However, due to the large genomes and relatively limited sample sizes, *trans-*eQTL remained difficult to detect in such settings, leaving part of the gene expression variance still unexplained^12^. Moreover, most population-level studies have focused on only a small fraction of genetic diversity, disregarding the wide variation in genetic content, namely the entire pangenome and more precisely the accessory genome. And finally, many species including humans display clear population structure that are often linked to the demographical, ecological and evolutionary histories of the subpopulations^17–19^. However, due to the sampling limitations, the impact of the subpopulation structure on gene expression remains unclear, and as a result, no unified view of the patterns of gene expression regulation within and among populations is currently available.

Understanding patterns of gene expression variation at a population-scale remains a challenge but it should provide deeper insights into the molecular basis of phenotypic diversity as well as transcriptional network architecture. Here, we took advantage of a population of 1,011 yeast isolates we previously completely sequenced to determine and explore the transcriptomic landscape at a population-scale^20^. The budding yeast *Saccharomyces cerevisiae* is a key model organism for investigating how genetic variants influence gene expression^6, 7, 21^. This species is characterized by a complex population structure with domesticated as well as wild subpopulations, and presents a high genetic diversity^20^. Large-scale genome analysis of the 1,011 natural isolates also provided a pangenome definition of the species as well a comprehensive view of genome variation at different levels, including copy number variants and gene content variation^20^. Finally, this large sample increases the power of genome-wide association studies (GWAS), allowing for in-depth characterization of local and distant regulatory variants impacting gene expression variation. Together, these two genomic and transcriptomic datasets have led to the most comprehensive insight into genome-wide expression regulation across the species, which would be a challenge to achieve at this scale and at the same level of accuracy for other organisms at present. Our study advances the understanding of the genetic and functional architecture of transcriptional landscape as well as its heritability at a species-wide scale.

**Figure 1.**
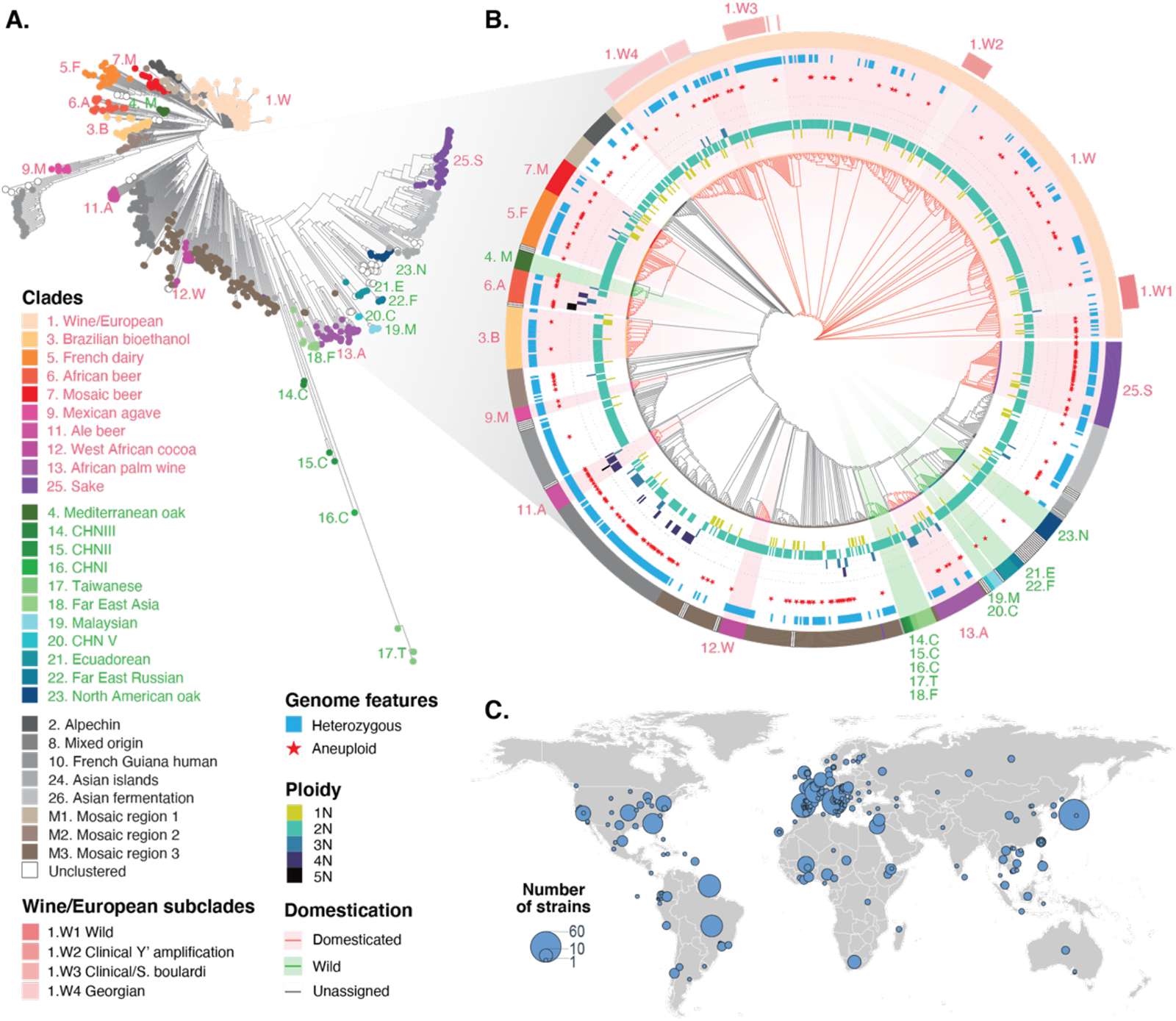
Origin and genomic diversity of 969 isolates. **A.** Neighbour-joining tree based on biallelic single nucleotide polymorphism among 969 isolates included in our data. Previously defined subpopulations^20^ are color-coded. **B.** Detailed descriptions of various genomic features for the isolates. From the inside to the outside: circular cladogram for the 969 isolates, coloured branches correspond to domestication (red) and wild (green) clusters; ploidy levels for each isolate ranging from 1N to 5N; presence of any aneuploidy marked as a star; heterozygosity marked as a blue bar; clades and subclades with indicated colour codes. **C.** Geographical distribution of the 969 isolates. The size of the circles indicates the number of isolates included from a given geographical location.

### Pan-transcriptome dataset generated

To gain a comprehensive overview of gene expression variation at the species level, we performed RNA sequencing for a collection of 1,032 *S. cerevisiae* natural isolates^20, 22^ and obtained 969 high quality transcriptomes with at least 1 million mapped reads (Figure 1, Table S1). The genomes of all these isolates have been previously completely sequenced and extensively characterized, reflecting the broad genetic diversity of the species in terms of single nucleotide polymorphisms (SNPs), gene copy number variants (CNVs), genome content variation (*e.g.*, introgressions, horizontal gene transfers) as well as aneuploidy and ploidy level variation (Figure 1A-B). The final set of 969 isolates are distributed across 26 well-defined clades that capture the ecological and geographical diversity of the species, including various domesticated and wild subpopulations (Figure 1A-C). Using the previously determined yeast pangenome^20^, we obtained the expression levels for 6,445 transcripts, including 4,977 core ORFs as well as 1,468 accessory ORFs that are variably present in isolates (Figure 2A, Table S2). This dataset constitutes the most comprehensive pan-transcriptomic catalogue within a single species to date and is essential to better understand what are the main drivers of gene expression variation, and consequently the key players that shape the transcriptional landscape in a population (Online Datafile 1).

### Accessory genes have unique transcriptional behavior

This population-scale dataset first allowed us to explore the transcriptional behavior of each of the 6,445 genes. Within this population, each gene can be characterized by its overall expression level as well as its variation across the 969 isolates. We therefore defined two metrics, namely “abundance”, which corresponds to the average expression level for a given gene across samples; and “dispersion”, which describes the variance across samples. The abundance is calculated as the mean log2 of the normalized read counts (transcript per million, or TPM) across isolates for which the gene is present in their genome. For the dispersion, we used the mean absolute deviation (MAD), which is more robust to outliers and do not assume normality of the expression levels compared to standard deviation (Methods).

By looking at the pangenome, we found that the core and accessory genes display distinct patterns, characterized by a significantly lower abundance and higher dispersion of accessory gene expression compared to core genes (Figure 2B-C). Beside this genomic view, we further investigated the transcriptional behavior of genes from a functional perspective. We examined 62 broad- and non-redundant biological process GO slim terms using gene set enrichment analyses (GSEA) based on rankings of the mean expression abundance and dispersion across all genes included in our dataset. Among the 62 GO slim terms, 59 are significantly enriched (FDR < 0.05) for abundance or dispersion, and are then grouped into three quadrants depending on the direction of the enrichments (Figure 2D, Table S3). Specifically, genes involved in GO terms related to growth and cellular metabolisms show high abundance and high dispersion. By contrast, genes involved in GO terms related to organelle organization, transcription regulation and protein homeostasis show high abundance but low dispersion. And finally, the low abundance and low dispersion quadrant is characteristic for genes involved in GO terms related to chromosome organization, DNA recombination and repair as well as cell cycle regulation (Figure 2D, Figure S1). Contrasting to accessory genes, none of the GO terms show a pattern of low abundance and high dispersion (Figure 2D, Figure S1), possibly suggesting that known biological processes that are lowly expressed tend to be more tightly regulated across different genetic backgrounds.

From a genomic and functional perspective, this survey of expression abundance and dispersion at the population level clearly highlights different transcriptional behaviors. The expression abundance alone is known to be correlated with different biological functions, such as the ribosomal genes are among the most highly expressed and genes involved in cell cycle regulation are in general lowly expressed and tightly regulated^23, 24^. Interestingly, expression dispersion, which is an added dimension here due to our large sample size, allows us to distinguish and identify sets of genes and pathways that are most likely to drive the transcriptional variation in a population. For example, genes involved in metabolism and growth are among the most highly abundant and dispersed biological processes, which could reflect the diverse metabolic states and preferences across different isolates. Remarkably, contrasting to genes in all known major biological processes, the accessory genes uniquely occupy the low-abundance and high-dispersion space and represent a previously under-characterized and unknown driver of transcriptional landscape diversity across the species.

**Figure 2.**
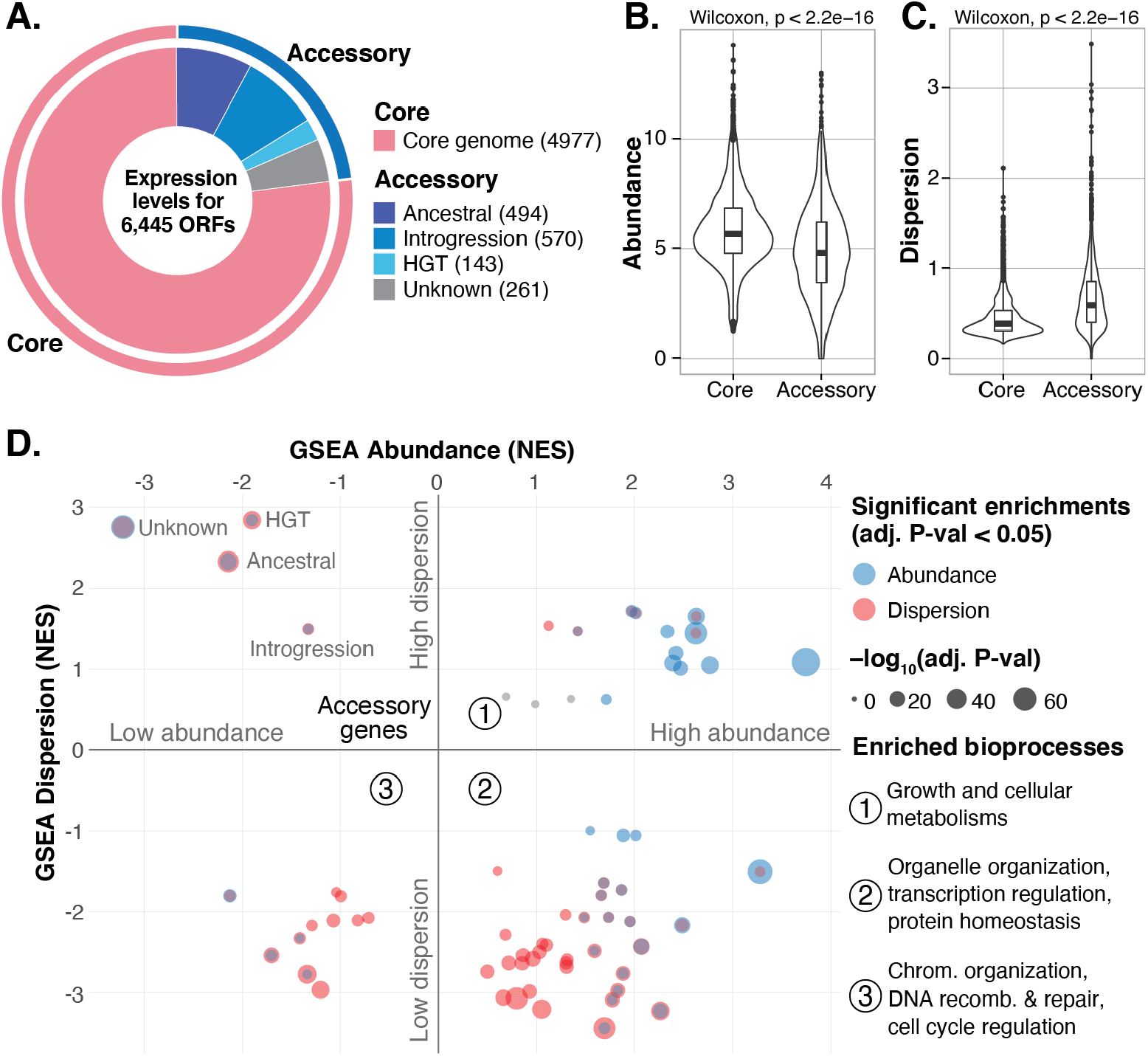
Functional description of the dataset. **A.** The number and distribution of all transcripts analysed in the data, including 4,977 core genes and 1,468 accessory genes as previously annotated based on the genomes^20^. **B-C.** Global comparison of mean gene expression abundance (**B**) and dispersion (**C**) between core and accessory genes. Mean expression abundance was calculated as the mean log2(TPM+1) across isolates and dispersion as the mean absolute deviation. For accessory genes, isolates that do not carry the given gene were excluded from the calculations. **D.** Gene set enrichment analyses results for expression abundance (y-axis) and dispersion (x-axis), presented as normalized enrichment scores (NES). A total of 62 non-redundant GO slim biological process terms and 4 accessory gene subcategories are included. Significant enrichments are coloured in blue (abundance) and red (dispersion). Summary terms for each quadrant are as indicated on the plot. Detailed distribution and enrichment for each term are presented in Figure S1.

### Co-expression patterns recapitulates the cellular functional network

Although expression abundance and dispersion provide relevant global transcriptional trends, they do not reflect the co-expression patterns among genes across the population. Exploring the gene co-expression network at the species-level is essential to better understand the coordination of gene expression of diverse cellular processes in genetically distinct isolates. This then makes it possible to investigate whether population structure can impact or shape such central wiring of cellular functions.

**Figure 3.**
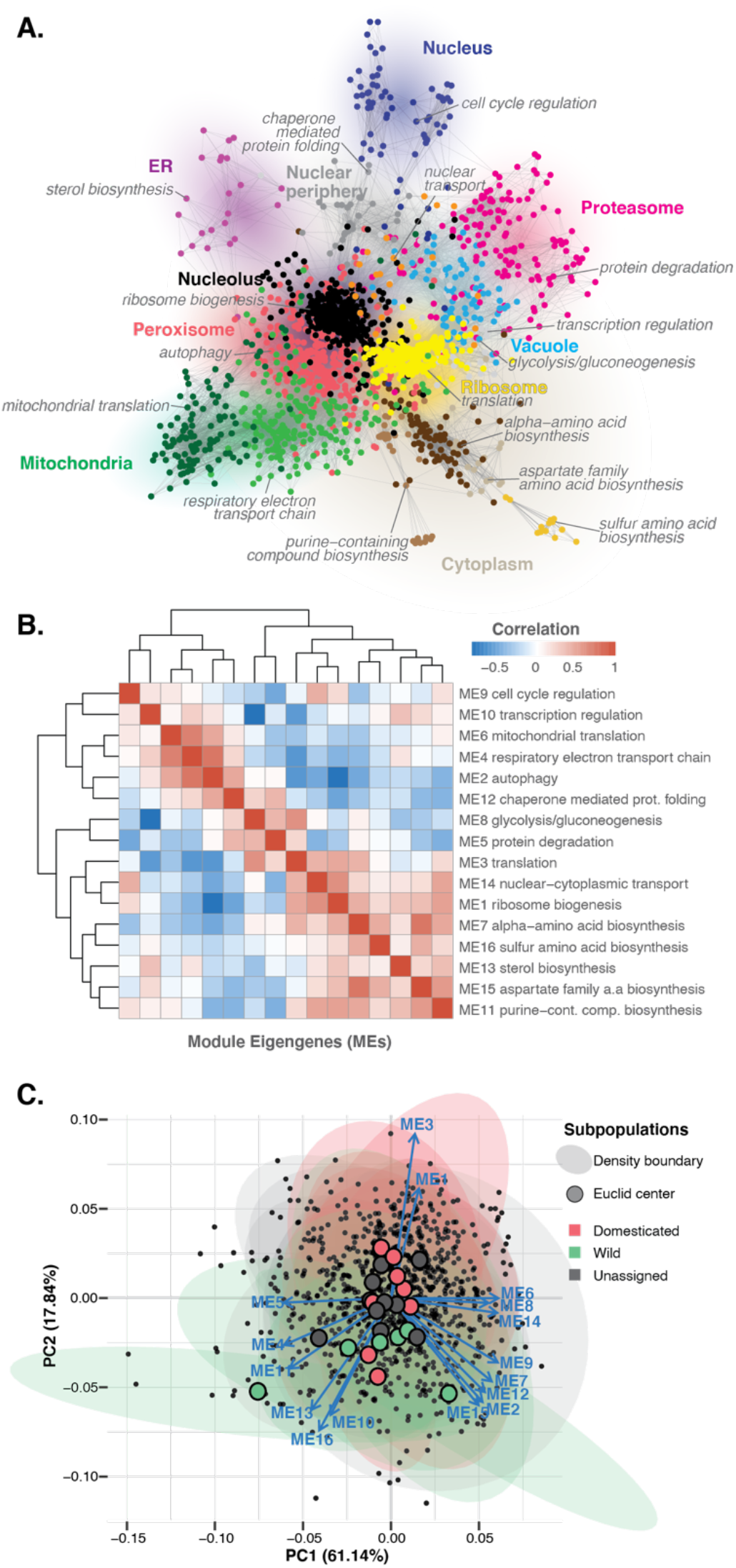
Whole population level gene co-expression network. **A.** Co-expression network based on pairwise gene expression profile similarities across 969 isolates. Nodes are coloured according to the 16 co-expression modules detected using weighted gene co-expression network analysis (WGCNA) method. The modules are annotated according to biological process GO term enrichment and are labelled in grey. Enriched cellular compartments are annotated and indicated in coloured shades. **B.** Pairwise similarity matrix based on eigengene expressions across the 16 modules. The modules are numbered according to descending order of their sizes. **C.** Principal component analysis (PCA) based on module eigengene expression across isolates. The first two principal components are plotted. The density boundaries are drawn for each subpopulation and are coloured according to the “domestication”, “wild” and “unassigned” clade annotations. The Euclid centres for each subpopulation-based density boundary are as indicated.

We therefore constructed a first species-wide gene co-expression network based on the pairwise expression profile similarities across the population (Figure 3A). Edges connect genes with similar expression profiles (Pearson’s r > 0.67) and genes with less than five edges were excluded. The resulting network consists of 1,797 genes displaying a scale-free architecture with a clear modular topology (Figure 3A). Using weighted correlation network analysis (WGCNA), we identified 16 co-expression modules localized to distinct regions of the network (Figure 3A, Table S4). Each module is enriched for a unique set of GO terms related to a similar biological process (Table S5), with the largest module (432 genes) enriched for *ribosome biogenesis*, and the smallest module (13 genes) corresponding to genes involved in *sulfur amino acid biosynthesis*. The relative positions of modules on the network also reflect broader functional relationship and shared cellular localizations (Figure 3A, Table S5). This hierarchical organization is further illustrated by examining the pairwise correlations between module eigengenes, which show that modules involved in distinct but related biological processes are clustered together (Figure 3B). Overall, these results clearly show that the defined co-expression patterns recap the cellular functional network.

Using module eigengene expressions, we performed principal component analysis (PCA) and focused on potential signatures related the subpopulation structures (Figure 3C). Globally, there is no clear delineation among different subpopulations based on the first two principal components (Figure 3C). This lack of strong subpopulation impact is corroborated by the general absence of differential subpopulation specific co-expression (Figure S2), indicating that the co-expression network is robust to genetic variation across the population.

Overall, this population-scale co-expression network captures the topological organization of the cell by displaying the hierarchical relationships between functionally defined modules. The network is globally robust to the population structure, highlighting the part of the transcriptional landscape that is coordinated, hierarchical and functionally conserved at the species level.

### Transcriptional signatures related to specific subpopulations and domestication processes

To identify subpopulation-specific transcriptional signatures, we performed differential gene expression analyses by comparing each clade to the rest of the population (Methods). We filtered out genes for which the expressions were not detected in over half of the population, resulting in an input set of 6,116 genes. We found 2,209 unique differentially expressed genes across clades (Table S4). The number of significant differentially expressed genes detected in a given clade does not necessarily correlate to the size of the subpopulation (Figure S3A, Table S4). On average, each subpopulation show ∼130 differentially expressed genes, ranging from 390 for the French dairy clade (5. F, 30 isolates) to 0 for the CHNII clade (15. C, 2 isolates) (Figure S3A, Table S4).

While the co-expression network reflects globally coordinated cellular processes at the population level, the differential expression set reveals variability in subpopulation-specific gene expression. To further characterize this aspect, we looked at the clustering of the individual isolates based on the expressions across the co-expression set (1,797 genes) and the differential expression set (2,209 genes), using t-distributed stochastic neighbor embedding (t-SNE) (Figure 4A-B). As expected, no structure related to the subpopulations is defined using t-SNE on the co-expression genes (Figure 4A). By contrast, a clear delineation of subpopulations, including multiple domesticated clades, can be observed using t-SNE on the differential expression genes (Figure 4B). Specifically, domesticated clades such as Wine/European (1. W), French dairy (5. F), African beer (6. A), Ale beer (11. A) and Sake (25. S), all show a clear and distinctive delineation, suggesting independent transcriptional signatures that are unique to each domestication process (Figure 4B). The mosaic subpopulations (M1 to M3 and unclustered) and the Brazilian bioethanol clade (3. B) show a more scattered pattern, which is consistent with their admixed genome structure. Interestingly, all the wild subpopulations, despite their high genetic divergence, show little transcriptomic differentiation and are all closely clustered together (Figure 4B). The West African cocoa (12. W) and the African palm wine (13. A) clades, although involved in human-related fermentative processes, are known to be derived directly from wild lineages and cluster more closely to the wild subpopulations (Figure 4C). The t-SNE clustering is further corroborated by the topologies of the neighbor-joining trees (Figure S4) based on Euclidean distances among isolates using either the co-expression (Figure S4B) or differential expression gene sets (Figure S4C). These observations show that the wild populations do not display differentiated expression patterns despite their high genetic divergence, and suggest that the differential expression landscape is mainly driven by multiple distinctive domestication processes.

**Figure 4.**
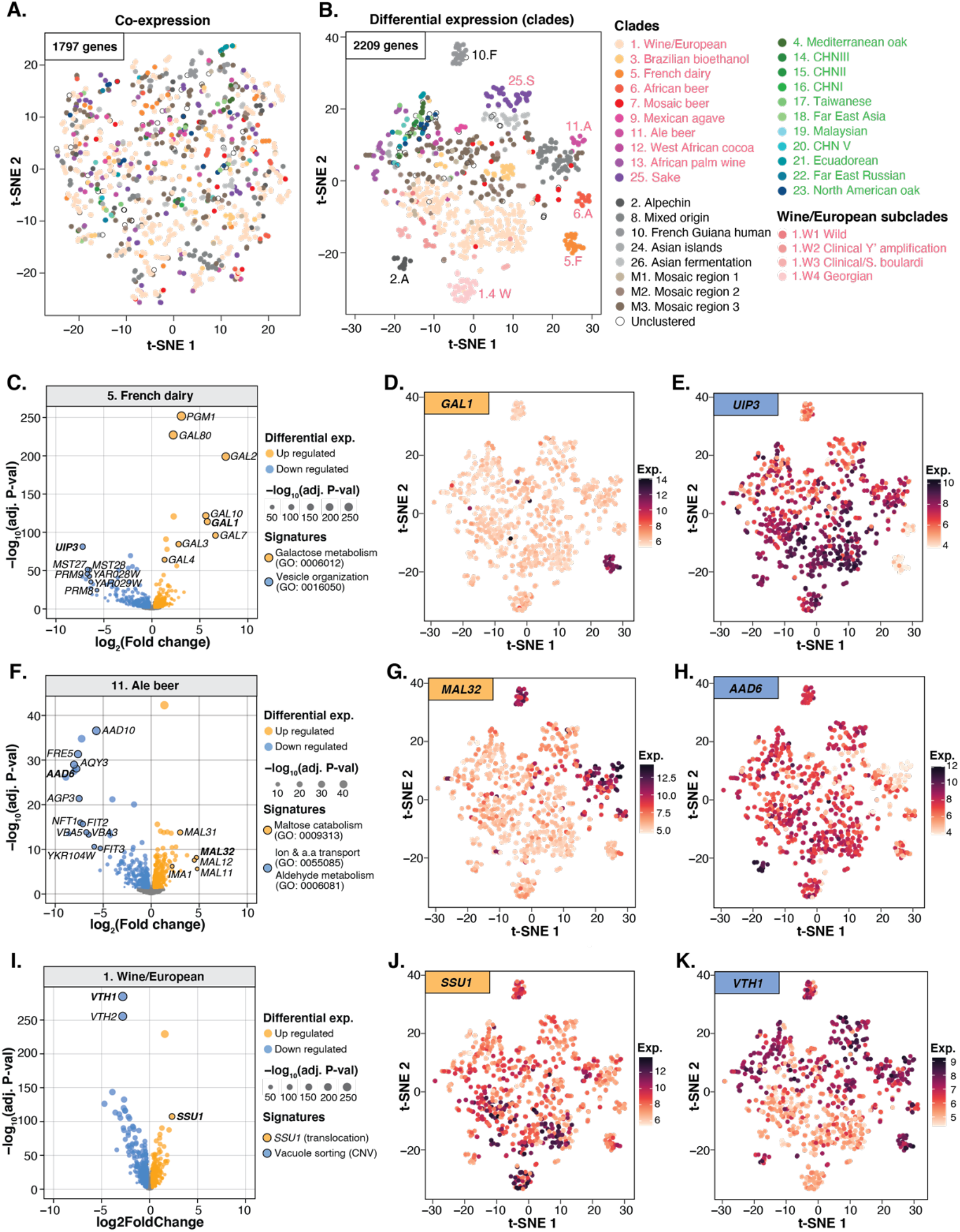
Subpopulation-specific differential gene expression. **A-B.** Isolate-level clustering using t-distributed stochastic neighbour embedding (t-SNE) based on the expressions of 1,797 co-expression genes (**A**) and 2,209 differential expression genes (**B**). Isolates belonging to each annotated subpopulation are colour coded. **C-K.** Examples of subpopulation-specific domestication signatures. Three examples are shown for signatures in 5. French dairy (**C-E**), 11. Ale beer (**F-H**) and 1. Wine/European (**I-K**). Volcano plots (**C, F & I**) present up-(orange) and down-regulated (blue) genes and biological processes, with x-axis showing the log2 fold-change comparing the subpopulation to the rest of the isolates, and y-axis showing the -log10 of Benjamini-Hochberg adjusted P-value (FDR). Dot sizes are scaled according to the FDR. The variance stabilized expression levels for specific up-(**D, G & J**) and down-regulated (**E, H & K**) genes are overlapped to the t-SNE plot, with the expression levels scaled as different colour intensities. The colour scales are included on the side of the plots.

To further characterize these transcriptional signatures, we performed GSEA based on the ranked log2 fold-change of the differential expression genes in each subpopulation (Figure S5, Table S8). Significant enrichments for various biological processes were found across different clades, the majority of which were known to be adaptive in specific domestication processes (Figure S3C). For example, genes in the *GAL* pathway, involved in the metabolism of galactose, are significantly up-regulated in the French dairy clade (5. F) (Figure 4C). In this subpopulation, the *GAL* pathway underwent from a tightly regulated glucose-repressed/galactose-induced system to constitutive expression even in the presence of glucose. Such switch was previously found in several lineages involved in spontaneous milk fermentation and was linked to adaptation to lactose rich medium^25, 26^. In addition to the *GAL* genes (Figure 4C-D), the French dairy clade also shows down-regulation of multiple putative integral membrane proteins in the *DUP240* family (*e.g.*, *MST27/28* and *UIP3*) that are involved in the COPI/COPII related vesicle organization^27^ (Figure 4C-E). Such changes in cell secretion could also be adaptive to certain cheese making processes^28^. In various types of alcohol fermentations, adaptive transcriptional signatures are also prevalent (Figure 4F-H, Figure S3C, Table S8). For example, the *MAL* genes involved in maltose catabolism are up-regulated in the Ale beer clade (11. A), a signature to malt fermentation environment in beer making^29^ (Figure 4F-G). At the same time, the expression of multiple aldehyde dehydrogenases genes (such as *AAD6* and *AAD10*) is down-regulated in the Ale beer cluster (Figure 4F-H). Similarly, down-regulation of another aldehyde dehydrogenase gene, *ADH7*, is seen in the Sake (25. S) cluster, along with a pathway-level up-regulation of genes involved in the thiamine metabolism (Figure S5, Table S8). Both these down- and up-regulation signatures are known to ensure high ethanol yield during sake production^30^. Another well-known adaptive trait in certain wine isolates was several translocations that lead to the overexpression of *SSU1*, a sulfite pump that confers resistance to sulfur dioxide, a commonly used compound in wine making^31, 32^. The overexpression of *SSU1* is indeed seen in the Wine/European clade (1. W) in our dataset (Figure 4I-K).

Differential expression analyses highlight transcriptional signatures that are specific to different subpopulations. Interestingly, wild subpopulations appeared to be less differentiated in terms of the transcriptional diversity despite the high level of genetic divergence among these clades. In contrast, domesticated subpopulations exhibit clear signatures that correspond to distinct adaptive processes, notably in various metabolic pathways uniquely selected in different domestication events.

### Introgression, horizontal gene transfers and gene expression variation

The recently established *S. cerevisiae* pangenome revealed numerous horizontally transmitted evolutionary events, such as introgressions and horizontal gene transfers (HGT), as part of the accessory genome^20^. This part of the pangenome is also more specific to certain subpopulations. For example, the presence of *Saccharomyces paradoxus* introgression events is a main characteristic of the Alpechin (2. A), Mexican agave (9. M) and French Guiana (10. F) clades^20^. By contrast, the Wine/European (1. W) clade features many HGT events coming from *Torulaspora* and *Zygosaccharomyces* species^20^.

Overall, the accessory genome is globally characterized by a low abundance and high dispersion of gene expression, as shown above. However, even though this is a strong general trend, a difference can be observed between the expression of introgressed genes and those originating from HGTs (Figure 2D, Figure 5A). In fact, while the gene expression abundance is higher, the gene expression dispersion is lower for introgressed genes compared to HGTs (Figure 5A). By looking at the gene expression level according to the species of origin, it is possible to explain the observed variation in terms of dispersion between introgression and HGT (Figure 5B). Regardless of the origin of the introgressed genes (*Saccharomyces paradoxus*, *Saccharomyces mikatae* or unknown *Saccharomyces* species), the level of gene expression is globally similar (Figure 5B). In contrast, there is a large disparity in gene expression level according to the species for the HGT events, leading to a higher dispersion (Figure 5B).

**Figure 5.**
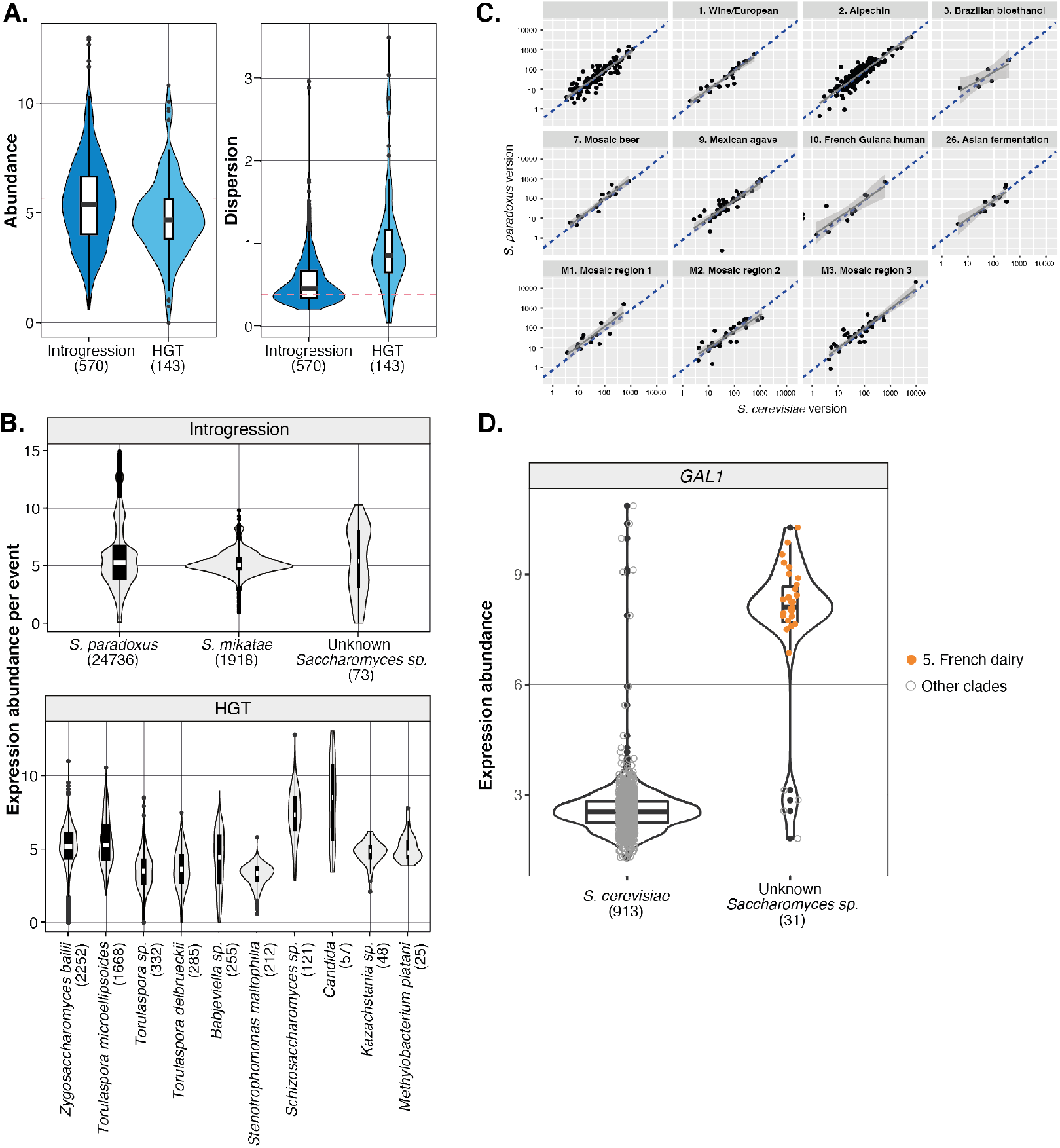
Global expression patterns for accessory gene subcategories. **A.** Mean expression abundance and dispersion for introgression (570) and horizontal transferred (HGT) genes (143). Dotted line indicates the median value observed in core genes. **B.** Expression abundance per event for the introgression and HGT subcategories by donor species. Expression abundance is calculated as log2(TPM+1). **C.** Expression correlation between the *S. paradoxus* and *S. cerevisiae* alleles in the introgression subcategory. Only heterozygous introgression events are presented, with the x-axis presenting the log10 scaled reads count for the *S. cerevisiae* version and the y-axis presenting the corresponding counts for the *S. paradoxus* version. Dotted lines indicate the one-to-one ratio. Panels correspond to subpopulations with large numbers of such introgression events. **D.** Expression abundance for the *GAL1* xenolog originated from an introgression event from an unknown *Saccharomyces* species. Expression abundance per isolate is presented on the y-axis and the origin of the *GAL1* alleles on the x-axis. Isolates from the 5. French dairy clade are highlighted.

The vast majority of introgressed ORFs examined here comes from *S. paradoxus*, a *S. cerevisiae* sister species. In most cases, these genes substitute their *S. cerevisiae* ortholog, either partially, resulting in a heterozygous state (one allele from each species), or completely, resulting in a homozygous state. To study the gene expression variation and adaptation of introgressed alleles, we first focused on the homozygous cases. We examined the expression of 437 homozygous genes for the allelic version of *S. cerevisiae* or *S. paradoxus* in different strains, and found that their expressions are well correlated (Figure 5C and Figure S6A). We next asked whether introgression events could impact expression level when they result in heterozygous states with *S. cerevisiae* native alleles. To do this, we performed Allele Specific Expression (ASE) analysis, minimizing the mapping bias towards the reference allele (Methods). Again, we found no significant difference in expression between *S. cerevisiae* and *S. paradoxus* versions, suggesting similar regulation of both alleles (Figure S6B). Overall, *S. paradoxus* introgressed alleles are therefore expressed at a level similar to those of *S. cerevisiae*, suggesting that they are well integrated in the transcriptional network.

Nevertheless, we also found some exceptions and a notable example corresponding to a gene whose donor species was identified as an unknown *Saccharomyces* species (Figure 5B). This set of alleles is characterized by the overexpression of the *GAL1* xenolog, involved in galactose metabolism. This allele is found in 31 isolates, 27 of which are dairy associated strains. In these strains, the *GAL1* gene is highly expressed compared to the non-dairy associated strains (Figure 5D). This gene was already identified in the differential expressed gene analysis as a signature of the French dairy clade (5. F). As mentioned previously, this is related to a rewiring of the *GAL* network with a constitutive expression of the *GAL* genes in the dairy related isolates^25^.

### Genetic basis underlying the pan-transcriptome variation

To further understand the relationship between the pangenome variation and the transcriptional landscape, we performed genome-wide association studies (GWAS) by considering both SNPs and CNVs that were previously characterized in the population^20^. Across the 969 isolates, 84,682 SNPs and 1,100 CNVs were included, with a minor allele frequency (MAF) higher than 5%. A total of 9,470 significant expression Quantitative Trait Loci (eQTL) were detected (Methods). In total, significant eQTL are associated with the expression variation of 3,471 genes. Among the detected eQTL, 7,273 are associated with SNPs and 2,197 are associated with CNVs, corresponding to 4,393 and 497 unique loci, respectively (Figure 6A-B, Datafile 2).

On the SNP-eQTL side, a total of 1,901 were found as local eQTL, with sites located in the upstream of the transcription start site (TSS) or within the open reading frame (ORF) of the target gene displaying the largest effect sizes (Figure S7A-B). The remaining 5,372 SNP-eQTL are distant and *trans*-acting. Overall, local SNP-eQTL are less frequent, representing ∼26% of the total set of eQTL detected, which is consistent with previous findings based on linkage mapping across a large segregant panel in a yeast biparental cross^6^. The *trans* eQTL detected are uniformly distributed across the genome, with ∼41% of eQTL (2,206 out of 5,372) impacting only one trait (Figure 6A). The most significant eQTL hotspot was mapped to the *CTT1* gene, encoding for a cytosolic catalase T, which is associated with 251 expression traits and was previously found to be an eQTL hotspot in stress conditions^33^ (Figure S7C). Contrasting to previous observations in a yeast cross^6^ and recent findings in a *C. elegans* population^5^, the set of *trans* eQTL detected in our large dataset are not biased toward a few hotspots with extreme pleiotropic effects (Figure S7C).

**Figure 6.**
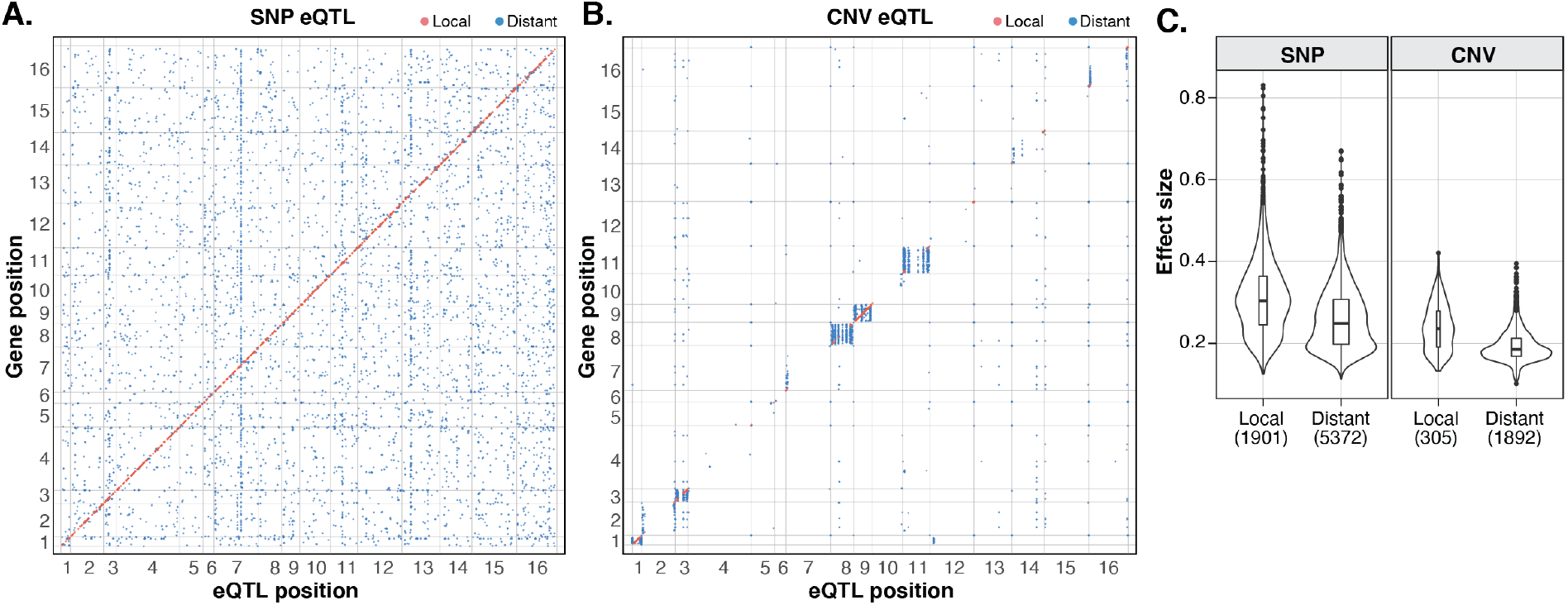
Genome-wide association identified SNP and CNV associated eQTL. **A.** Locations of local and distance SNP eQTL along the genome. SNP variant associated with an expression trait are on the x-axis and the position of the genes are on the y-axis. Local eQTL are coloured in red and distant eQTL in blue. **B.** Locations of local and distant CNV eQTL. Local eQTL correspond to associations between the CNV and expression trait of the same gene and are coloured in red. Distant eQTL are coloured in solid blue. **C.** Comparison of the absolute effect sizes for local and distant eQTL for SNP and CNV eQTL types. Significant differences are indicated with stars. See also Figure S7.

Contrasting to SNPs, the effect of CNVs on gene expression variation has never been systematically explored at the species scale. Compared to SNPs, CNVs are not randomly distributed along the genome and tends to be located towards the subtelomeric regions, except for chromosomes 1 and 9 due to the presence of aneuploidies that passed the 5% MAF filter (Figure 6B). In addition, chromosomes 3, 8 and 11 are also impacted by aneuploidies at a lower frequency (∼3%), resulting in a larger number of CNV eQTL in those regions (Figure S7D). These aneuploid chromosomes artificially inflates the *trans* hotspots for CNV-eQTL (Figure S7D). We only considered CNV-eQTL to be local when the CNV is directly associated with the same gene expression trait, and distant CNV-eQTL that correspond to single linkage groups. This results in a total of 305 local CNV-eQTL *versus* 1,892 distant CNV-eQTL. On average, each CNV-eQTL impacts about eight expression traits.

Consistent with previous observations, local SNP-eQTL display larger effect sizes compared to the distant ones, with a 1.3-fold higher absolute effect sizes and 2.4-fold higher variance explained on average (Wilcoxon p-value <2.2e-16) (Figure 6C, Figure S7E). While the same trend holds true for CNV-eQTL for absolute effect sizes (1.2-fold higher absolute effect sizes, local vs. distant, Wilcoxon p-value <2.2e-16) (Figure 6C), the variance explained by local or distant CNV-eQTL are low and not significantly different due to an overall lower minor allele frequency of CNVs compared to SNPs (Figure S7E). Overall, CNV-eQTL display smaller effect sizes compared to SNP-eQTL across the board. This first direct comparison of eQTL effect sizes indicate that SNPs display significantly larger impact than CNVs for gene expression variation at the population level.

From a functional perspective, *trans* eQTL uncovered coherent associations that link causal SNPs and gene expression traits within the same biological process (Figure S7F). The top 5 *trans* eQTL hotspots collectively impact the expression of 356 genes, of which 276 belongs to the *ribosome biogenesis* module on the co-expression network (Figure S7F). The causal SNPs mapped to *CTT1* (251 eQTL, chromosome 7, position 655,851), *SRD1* (84 eQTL, chromosome 3, position 148,921), *DHR2* (82 eQTL, chromosome 11, position 290,740), *RAD52* (71 eQTL, chromosome 13, position 213,896) and *RPS17A* (62 eQTL, chromosome 13, position 225,572) (Figure 6B), of which *SRD1* and *DHR2* are involved in rRNA processing and synthesis, and *RPS17A* is a ribosomal protein, all of which are directly related to ribosomal biogenesis. Furthermore, *trans* eQTL hotspots appeared to affect disproportionally more genes on the co-expression network. Among the eQTL hotspots that are associated with more than 20 traits, 648 gene expressions are affected and of which 404 belongs to the co-expression network *vs.* 105 that belong to the differential expression genes (Table S9).

By integrating the eQTL results with the global transcriptome structure, we uncovered distinctive patterns regarding the genetic basis underlying the co-expression and differential expression genes (Figure S8A-C). Overall, differential expression genes are significantly more likely to be controlled by any eQTL compared to the total set (Odds ratio 1.1, two-sided Fisher test p-value = 0.01) (Figure S8A), while co-expression genes are slightly depleted (Odds ratio 0.92, Fisher test p-value = 0.08). However, the types of eQTL involved showed more drastic differences, with a 0.38-fold depletion of local eQTL (two-sided Fisher test p-value < 2.2e-16) for co-expression genes and a 1.42-fold enrichment of local eQTL in the differential expression genes (two-sided Fisher test p-value = 1.921e-08) (Figure S8B). CNV-eQTL are also significantly depleted in co-expression genes (Odds ratio = 0.62, two-sided Fisher test p-value = 4.553e-07) (Figure S8C).

**Figure 7.**
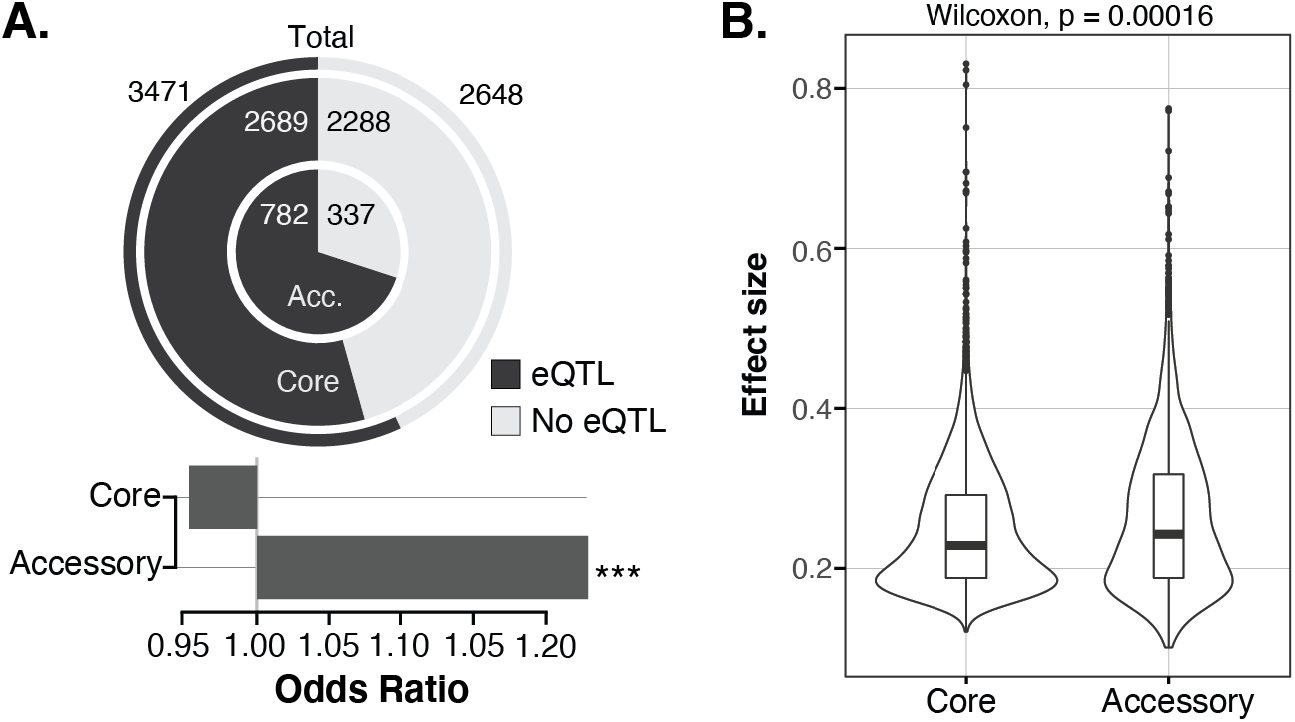
Accessory genes show proportionally more eQTL with higher effect sizes. **A.** The number and proportion of genes that are impacted by at least one eQTL for accessory genes (inner ring), core genes (middle ring) and the combined set (outer ring). Fold-enrichment for core and accessory genes on the proportion of genes impacted compared to the combined set are presented as bars (odds ratio). Significant enrichment based on the two-sided Fisher test is indicated with stars. **B.** Comparison of the absolute effect size for eQTL associated with accessory or core gene expression traits. See also Figure S8.

From the perspective of the pangenome, accessory genes are significantly more likely to be controlled by at least one eQTL compared to the core genes (Figure 7A). Accessory genes are also significantly more likely to be controlled by local eQTL (Odds ratio = 1.33, two-sided Fisher test p-value = 0.0002676) (Figure S8D-E). Most remarkably, the effect size for eQTL associated with accessory genes is globally higher compared to core genes (Figure 7B), and the same trend holds true for the fraction of variance explained (Figure S8F). These differences are not biased toward accessory genes with a low occurrence in the population (Figure S9). Overall, these observations clearly show that the accessory genome is a key component of gene expression variation regulation at a population-scale.

## Discussion

The species-wide pan-transcriptomic analysis presented here has led to a precise characterization of the functional organization and the genetic basis underlying the transcriptional landscape at a scale that is not yet achieved in any other species. Our results revealed the accessory genome as a key driver of the transcriptional diversity, contributing proportionally more to heritable expression variation than the core genome.

Natural population of *S. cerevisiae* is highly diverse and displays clear population structure, with defined subpopulations assigned to specific domesticated and wild lineages^25, 34, 35^. Such population structure is commonly observed in various species, including humans^18^, however, the impact of population structure on the transcriptional landscape remains largely unclear. We characterized gene expression patterns both at the whole population level using co-expression analysis, and at the subpopulation level using differential expression analysis. Our results show that the global transcriptional landscape is consistent with a two-tier architecture, characterized by a tightly interconnected central network (*i.e.* co-expression) and an auxiliary structure related to differentiated gene expression patterns (*i.e.* differential expression). These two architectural levels are not equally impacted by the population structure. On one hand, the co-expression network captures the main biological functions, reflects the topological organization of the cell and is globally conserved across the subpopulations. On the other hand, differential expression reveals subpopulation-specific, functionally coherent up- and down-regulations that can be associated with distinct domestication signatures.

From the pangenomic perspective, the accessory genome, including ancestrally segregating genes in the *S. cerevisiae* species and horizontally acquired genes from close (introgression) or distant (HGT) relatives, all exhibit higher expression dynamics compared to the core genome. The core and accessory gene features roughly echo the two-tier transcriptomic landscape, with accessory genes more likely to be involved in the auxiliary network than in the central network. In addition, expression patterns suggest that the accessory genes, despite being variable across the population, can also be important to certain adaptive processes and represent an integral part of the functional genome.

The large population size and the fully catalogued genetic variants allowed us to systematically explore the genetic basis underlying transcriptional variation at the species level. By performing genome-wide association studies with both SNPs and CNVs, we uncovered local and distant eQTL for over 56% of the expression traits, with SNP-eQTL explaining a significantly higher fraction of variance compared to CNV-eQTL. Overall, local eQTL explain a higher fraction of variance, which is consistent with previous observations^5, 6^. However, distant eQTL are mostly randomly distributed along the genome and are not biased toward a few extreme hotspots, unlike the previous observation in a single cross^6^. Considering that these hotspots were possibly related to large effect rare variants with extreme pleiotropy, it is not surprising that such a pattern is not conserved in a large natural population where the genomic constitution is much more complex. Finally, accessory genes are significantly more likely to be associated with eQTL than the core genes. Moreover, eQTL associated with accessory genes also explain a higher fraction of the expression variance.

Taken together, our analyses at all levels collectively show the surprisingly high impact of the accessory genes on the transcriptional landscape within a species. While the accessory genome likely explains some of the missing heritability, the understanding of genetic effects on cellular phenotypes is far from complete. Dissecting the genetic regulation of an additional set of molecular phenotypes or intermediates, such as proteomes, will most likely yield additional insights.

## Methods

### Description of the isolates and sample preparation

A collection of 1,032 isolates was compiled from the 1,011 strains collection^20^ and ^22^, along with the lab reference strain S288C (FY4-6). We measured the growth for all strains using 96-well liquid growth in standard synthetic complete (SC) media with 2% glucose as the carbon source. The growth rates were then extracted based on continuous OD measurements during 48 hours at 30°C in a microplate reader (Tecan Infinite F200 Pro). The strains were reorganized and grouped in 96-well plates according to their growth rates and then grown in 1 ml of liquid SC media using deep well blocks until reaching the mid-log phase (OD ∼0.3). For each sample, 750 µl of the culture was collected and then transferred to a sterile 0.45µM 96-well filter plates (Norgen, #40008) on a vacuum manifold (VWR, #16003-836). We applied vacuum to remove all the SC medium, sealed with aluminium foil seals, and flash froze the entire plate in liquid N2 to store the plate at -80°C before mRNA extraction. A final set of 969 isolates were included in our dataset after controlling for the final OD reading at the culture collection step as well as the quality of the RNA sequencing data. Detailed description of the isolates can be found in Table S1.

### RNA extraction, library preparation and sequencing

For each filter plates, mRNA was extracted using the Dynabeads® mRNA Direct Kit (ThermoFisher, #61012) based on an optimized protocol for high-throughput RNA sequencing^6^. Cells were lysed using glass beads and lysis buffer, then incubated for 2 minutes at 65°C. After RNA precipitation, two rounds of cleaning were performed using magnetic beads coupled to oligo (dT)_25_ residues which can hybridize to the polyA tails of the mRNA. A final volume of 10 µl of purified mRNA was obtained to prepare the sequencing library.

Sequencing libraries were prepared with the NEBNext® Ultra™ II Directional RNA Library Prep Kit for Illumina (NEB, #E7765L) in 96-well plates. We used 5 µl of purified mRNA for the library prep, corresponding to ∼10 ng of RNA molecules per sample. We generated cDNA libraries using reverse transcription. The resulting cDNA libraries were then purified using NEBNext sample purification magnetic beads and eluted in 50µl of 0.1X Tris-EDTA buffer. Dual index duplex adapters were added to the cDNA by ligation. In total, 96 combinations of TS HT dual index duplex mixed adapters from IDT® (Integrated DNA Technologies®) were used and each prepped DNA was assembled to a unique barcode combination. The adaptor ligated cDNA was purified using NEBNext sample purification magnetic beads and eluted in 15 µl of 0.1X Tris-EDTA buffer. A final PCR enrichment of the barcoded DNA were performed in a 9-cycle amplification using Illumina P5 and P7 universal primers (P5 IDT: 5’-AATGATACGGCGACCACCGA-3’; P7 IDT: 5’-CAAGCAGAAGACGGCATACGA-3’). 21 µl of final barcoded DNA were purified and eluted in 0.1X Tris-EDTA buffer.

For each sample, the final barcoded DNA was quantified using the Qubit™ dsDNA HS Assay Kit (Invitrogen™) in 96-well plate with a microplate reader (Tecan Infinite F200 Pro), excitation laser set at 485nm and emission laser at 528nm. All the samples from the 96-well plate with a concentration higher than 1 ng/µl were grouped and 20 ng of cDNA were collected and pooled from each sample. The DNA integrity of the pool was controlled on 1% agarose gel and quantified on Nanodrop and Qubit using the Qubit™ dsDNA HS Assay Kit (Invitrogen™).

The final pool of DNA was sequenced on Nextseq 550 high-output at the EMBL Genomics Core Facilities. In total, 1,046 samples were sequenced, including duplicates for some of the isolates. On average, 6.45 millions of 75 bp single-end reads was obtained for each sample after demultiplexing (Table S1).

### Reads cleaning and data processing

Raw reads were cleaned with cutadapt^36^ to remove adapters as well as low quality reads that were trimmed on the basis of a Phred score threshold of 30 and discarded if less than 40 nt long after this trimming step.

For each of the 1,046 samples, clean reads were mapped to the *S. cerevisiae* reference sequence using TopHat (v2.0.13)^37^. The resulting bam files were sorted and indexed using SAMtools (v1.9)^38^. Duplicated reads were marked using Picard (v2.18.14) in GATK (v4.1.0.0)^39^. HaplotypeCaller was used to call variants in each individual sample. The variant calling files (VCFs) were merged and the rare SNPs, defined as having a Minor Allele Frequency (MAF) less than 5% were extracted and intersected with SNP data from Peter et al. 2018 using bcftools isec.

The 1,046 samples were ranked based on the number of shared rare SNPs with each relevant strain described in the SNP matrix. This allows to automatically validate 940 unique isolates for which the expected strain was among the top 3 ranking strains. The remaining samples were manually investigated: 24 samples that were part of a large cluster of closely related strains could be validated as the expected strain and 19 samples could be unambiguously reassigned to the top 1 ranking strain. 14 samples out of the 1,046 could not be validated or reassigned and were discarded from the remaining analyses. After this step, a final set of 987 unique isolates was retained.

### Gene expression quantification

For each validated sample, reads were mapped to the *S. cerevisiae* reference sequence in which the SNPs of the corresponding strains were inferred (as described in Peter et al. 2018) plus the accessory genes that were not classified as ancestral or *S. paradoxus* orthologs in Peter et al. 2018 (n = 395). The mapping was achieved using STAR^40^ with the following parameters:

--outReadsUnmapped Fastx \

--outSAMtype BAM SortedByCoordinate \

--outFilterType BySJout \

--outFilterMultimapNmax 20 \

--outFilterMismatchNmax 4 \

--alignIntronMin 20 \

--alignIntronMax 2000 \

--alignSJoverhangMin 8 \

--alignSJDBoverhangMin 1

Isolates with more than 1 million reads mapped were kept for analysis, resulting in a final set constituted of 969 strains (Table S1).

Mapped reads counts were then obtained using the featureCounts function from the Subread package^41^ with the genes described in the *S. cerevisiae* reference annotation (n = 6,285) and accessory genes (n = 395) as features. The following options were used in order to get multi-mapped reads counted as a fraction of the sites they mapped to:

-M \

--fraction

Finally, Transcripts Per Million (TPM) were calculated as a measure of transcription abundance for each of those features and a log2(TPM+1) transformation was applied. From the set of 6,285 reference genes, 196 were filtered out because log2(TPM+1) was lower than 1 in 50% of the isolates. The read counts for 39 accessory features with a known homolog in *S. cerevisiae* according to the pangenome annotations were merged with the corresponding homolog. The final set is thus constituted of 6,445 ORFs which were used for downstream analyses (Table S2).

### Neighbor joining tree

The variant calling files of the 969 final strains were combined using GenotypeGVCFs in GATK. The biallelic segregating sites were used to construct a neighbour-joining tree with the R packages ape^42^ and SNPrelate^43^. Briefly, the .gvcf matrix was converted into a .gds file for individual dissimilarities to be estimated for each pair of individuals with the snpgdsDiss function. The bionj algorithm was then run on the obtained distance matrix.

### Calculating mean expression abundance and dispersion

In total, 944 out of the 969 isolates in the final dataset were present in 1,011 isolates characterized previously^20^. For these isolates, the pangenome annotations in terms of the presence and absence of a given gene in each isolate are available. We used this set of isolates and their expression levels to calculate the mean expression abundance and dispersion. The abundance corresponds to the mean expression levels for all isolates where the gene is annotated as present. The dispersion is calculated as the mean absolute deviation using the following formula:

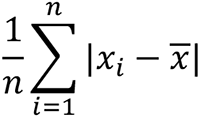

Where *n* is the number of strains that carries the gene, *x_i_* is the expression level in log2(TPM+1) for the *i*th isolate and 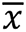 is the mean log2(TPM+1) for all samples for a given gene. Genes that are present in only one isolate were excluded. Genes that are not expressed in any isolates were also excluded. In total, 6,138 genes passed the filters and were included in the analysis, including 1,291 accessory and 4,847 core genes. All annotations can be found in Datafile S1.

### Variance stabilizing transformation

We performed variance stabilizing transformation using raw counts for each gene across the 969 isolates. We excluded genes that were not expressed in over half of the samples, which eliminated the majority of accessory genes originated from HGT. The remaining 6,119 genes were normalized using the vst() function in the R package DEseq2^44^. The variance stabilized expression values were subsequently used for co-expression and differential expression analyses.

### Gene co-expression analysis and module detection

We calculated Pearson’s correlation between all pairwise combinations in the 6,119 variance stabilized gene expressions. We generated an adjacency matrix by removing any gene pairs with an absolute correlation coefficient less than 0.67, then created an undirected network graph using the igraph package in R. We calculated the connectivity for each node and recursively removed nodes that are connected by less than 5 edges. This resulted in a final graph that contains 1,797 nodes and 181,954 edges. Graphic representation of the network was calculated using the fruchterman-reingold layout in the sna package in R^45^.

To detect co-expression modules, variance stabilized expression for the 1,797 node genes were used to generate Topological Overlap Matrix (TOM) using the R package WGCNA^46^. We performed scale independence test and determined the soft-threshold beta value, also known as the power value. At a beta of 5, the scale-free topology model fit was stabilized at a R^2^ of 0.9. The TOM is therefore calculated based on the signed adjacency matrix with the power of 5, using the TOMsimilarity() function in WGCNA. The dissimilarity matrix is then calculated as 1-TOM.

A distance matrix based on the TOM dissimilarity was calculated using the as.dist() function. Clustering was performed using hclust() in the fastcluster package, with the “average” method^47^. We used the cutreeDynamic() function in the dynamicTreeCut package^48^ to determine topologically independent clusters, with the options cutHeight = 0.95 and minClusterSize = 5. These clusters were treated as pre-modules, for which the module eigengene expressions were determined using WGCNA^46^. These pre-module eigengene expressions were clustered based on the dissimilarity of the pairwise correlation matrix, again using hclust() with method = “average”. Based on this, eigengenes with a dissimilarity < 0.2 were merged, forming a final set of 16 modules (Table S4). GO term enrichment analyses were performed using annotations for both the biological process (BP) and the cellular compartment (CC) standards^49^, using the mod_ora() function in the CEMiTool package^50^. Detailed enrichment results are in Table S5.

For the final 16 modules, eigengene expression was calculated using the function moduleEigengenes() in WGCNA. Principle component analysis (PCA) based on the eigengene expressions were performed using the prcomp() function in the stats package.

### Subpopulation specific differential expression analyses

The variance stabilized expression dataset comprised of 6,116 genes was used to perform subpopulation specific differential expression analyses using DEseq2^44^. Each subpopulation was compared to the remaining isolates using the expression matrix and annotated isolate information. Around 10% of the isolates in our dataset were haploids with defined mating types. To eliminated the effect of mating type specific expression signature, we incorporated the mating type information in the design model as a covariate. Significant differential expressions were determined using the Benjamini-Hochberg adjusted P-value less than 5%, corresponding to a 5% false discovery rate (FDR).

Due to the imbalanced subpopulation sizes, the FDR cutoff alone is biased toward detecting more differential expression with small effect sized in larger subpopulations. To remove this bias, we repeated the analyses by setting a cutoff on the absolute log2 fold-change ranging between 0 to 1 with an increment of 0.05. We counted the number of significant hits based on each cutoff and evaluated its relationship with the subpopulation size. We found that the dependency between the number of significant hits and the subpopulation size was removed around a cutoff of absolute log2 fold-change of 0.2 to 0.3. We therefore chose absolute log2 fold-change > 0.3 and adjusted P-value less than 5% as the final criteria for significant hits. All significant differential expression hits are presented in Table S6. Hits that overlapped with the co-expression genes were mainly associated with the Ecuadorean (21. E) and the French Guiana human (10. F) subpopulations, and were not included in the differential expression gene set for subsequent analyses.

### Gene set enrichment analyses

Gene set enrichment analyses (GSEA) were performed for gene-level abundance and dispersion analyses (Figure S1), co-expression module over-representation (Figure S3B) and for differential expression sets (Figure S3C).

GSEA on expression abundance/dispersion was performed using the fgsea R package^51^. Annotation of the GO slim terms was obtained from the SGD database. To reduce redundancy, we used the rrvgo R package and calculated the similarities between each annotated term in a pairwise manner using the “Resnik” method, and removed terms that are at least 70% overlapping with another term. We then performed GSEA using the fgsea() function, the pathways corresponding to the reduced GO slim terms and a size limit on the terms between 50 and 600. In total, 100,000 permulations were performed. The results are found in Table S3.

We performed GSEA based module over-respresentation test using the mod_gsea() function in CEMiTools^50^, to test for subpopulation related differential co-expression. Subpopulations with less than 10 isolates were removed to ensure statistical power. The results are found in Table S7.

Finally, GSEA was performed to identify GO term enrichments for subpopulation-specific differential expressions. For each subpopulation, significant hits were ranked by the log2 fold-change and then tested for enrichment in standard GO terms for annotated biological processes (BP), with term size limits between 5 and 500. For each test 10,000 permutations were performed. The results are shown in Table S8.

### Allele specific expression (ASE)

We selected all the isolates previously described as diploid, euploid and heterozygous^20^ in order to perform ASE analysis on this population (n = 289). We quantified the biallelic expression of each of these isolates using the GATK tool ASEReadcounter^52^ by providing it for each isolate a BAM file resulting from an alignment of RNA-seq read on the reference genome and a VCF file containing all heterozygous positions of the corresponding isolate. Heterozygous sites displaying a risk of allelic mapping bias were detected using their 75 bp mappability from GenMap^53^. We used the allele count to calculate an alternative allele ratio (AAR):

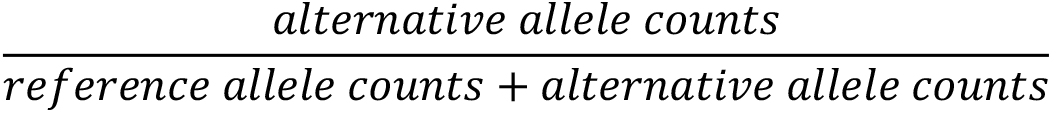

We finally excluded sites which did not have their heterozygosity supported by their alternative allele ratio (AAR = 0 or 1).

We detected imbalance in the allele expression using a simple binomial test corrected by FDR. In order to further compensate residual mapping bias in our results, we set the probability value of the binomial test to the mean of the alternative allele ratio in all our 289 isolates instead of 0.5. Moreover, we performed the previous test on sites that were covered more than 29X in order to ensure enough statistical power to our binomial test. Finally, we limited our explorations of ASE to the heterozygous sites located in CDS. In total, a list of 214,551 heterozygous sites distributed in 3,570 unique gene was analyzed across our 289 isolates (median = 464 sites per isolate.

Besides homozygous *S. paradoxus* introgression, heterozygous cases of *S. cerevisiae* and *S. paradoxus* alleles were also identified in the species. The unfiltered VCF files from Peter et al. 2018 were corrected for coverage and mapping bias, allowing to get 3,338 sites related to heterozygous introgressed genes. A significant difference was found in terms of AAR between these 3,338 introgressed sites and the non-introgressed toward low values for introgression. However, among those sites, some were displaying aberrant genetic allele balance (AB tag in the VCF file) due soft filtration. Thus, we iteratively performed several filtration steps of the genetic allele balance. In brief, at each step, the filtration value was set to exclude extreme genetic allele balance: for example, with a filtration value of 0.1, the site with a genetic allele balance higher than 0.9 or lower than 0.1 were discarded, for 0.2 the threshold was 0.8 and 0.2. Ultimately, this led to selecting sites with genetic allele balance narrowed to 0.5 but also resulted in an important decrease in the number of sites. In addition, at each filtration step, we compared the AAR between heterozygous introgressed sites and non-introgressed sites and found that the AAR difference between introgression related sites and the other sites decreased as the filtration value increased. Because extreme genetic allele balance could be related to difference in term of allele copy number and since our goal was to compare the allele balance in gene with similar genetic organization in their allele, we finally selected sites with a genetic allele balance between 0.33 and 0.66. This resulted in 356 sites distributed in 43 heterozygous introgressed genes.

### Genome-wide association studies (GWAS)

Genome-wide association studies based on mixed-model association analysis were performed as described in Peter et al., 2018 using FaST-LMM^54^. For SNP-eQTL detections, we removed the sub-telomeric regions corresponding to 20kb each side of the chromosomes from the SNPs matrix. A total of 84,682 SNP sites, and 1,100 CNVs between 969 strains with a minor allele frequency (MAF) higher than 5% were tested. For CNV-eQTL, as most variants were located in the subtelomeric regions, all expression traits were included. In total, the expression variation (in TPM) of 6,119 genes were tested. The SNP matrix was used for kinship for both SNP and CNV GWAS in order to account for population structure. Trait-specific p-value threshold was established for each gene by permuting phenotypic values between individuals 100 times. The significance threshold was the 5% quantile (the 5^th^ lowest p-value from the permutations) in each set, then Benjamini-Hochberg adjusted to account for multiple test bias. The effect size and variance explained by each variant was computed with FaST-LMM, with the effect sizes corresponding to the absolute value of “SnpWeight”, and variance explained corresponding to “EffectSize” from the raw output. The significance thresholds were scaled to account for the different sizes of the SNP and CNV matrices. Genomic inflation factor was calculated for each trait and the P-value was corrected when the genomic inflation factor is higher than To account for linkage disequilibrium among SNP and/or CNV loci, we grouped significant variants with an R2 > 0.6 and collided them into a single linkage group. Within each associated linkage group, the variant with the most significant association was kept. Prior to filtering, 12,058 SNP-eQTL and 47082 CNV-eQTL were detected as significant. After colliding the linkage groups, these numbers were reduced to 7,273 and 2,197 for SNP- and CNV-eQTL, respectively. For SNP-eQTL, local and distant eQTL were distinguished according to the distance from the considered gene: local eQTL can be located 25 kb each side around the gene, all other being considered as distant eQTL. For CNV-eQTL, we considered them local only when the variant and the associated trait are the same ORF. Significant associations can be found in Datafile S2 and are summarized in Table S9.

## Data availability

All sequencing reads are available in the European Nucleotide Archive (ENA) under the accession number PRJEB52153.

The 1002 Yeast Genome website – http://1002genomes.u-strasbg.fr/files/ – (RNAseq) provides access to:

- Datafile S1: final_data_annotated_merged_04052022.tab
- Datafile S2: GWAS_combined_lgcCorr_ldPruned_noBonferroni_20221207.tab

## Code availability

All custom scripts used in this study are available upon request.

## References

1. Hill, M.S., Vande Zande, P., and Wittkopp, P.J. (2021). Molecular and evolutionary processes generating variation in gene expression. Nat. Rev. Genet. 22, 203–215. 10.1038/s41576-020-00304-w.

2. Albert, F.W., and Kruglyak, L. (2015). The role of regulatory variation in complex traits and disease. Nat. Rev. Genet. 16, 197–212. 10.1038/nrg3891.

3. Schadt, E.E., Lamb, J., Yang, X., Zhu, J., Edwards, S., GuhaThakurta, D., Sieberts, S.K., Monks, S., Reitman, M., Zhang, C., et al. (2005). An integrative genomics approach to infer causal associations between gene expression and disease. Nat. Genet. 37, 710–717. 10.1038/ng1589.

4. Vande Zande, P., Hill, M.S., and Wittkopp, P.J. (2022). Pleiotropic effects of trans-regulatory mutations on fitness and gene expression. Science 377, 105–109. 10.1126/science.abj7185.

5. Zhang, G., Roberto, N.M., Lee, D., Hahnel, S.R., and Andersen, E.C. (2022). The impact of species-wide gene expression variation on Caenorhabditis elegans complex traits. Nat. Commun. 13, 3462. 10.1038/s41467-022-31208-4.

6. Albert, F.W., Bloom, J.S., Siegel, J., Day, L., and Kruglyak, L. (2018). Genetics of trans-regulatory variation in gene expression. eLife 7, e35471. 10.7554/eLife.35471.

7. Kita, R., Venkataram, S., Zhou, Y., and Fraser, H.B. (2017). High-resolution mapping of cis – regulatory variation in budding yeast. Proc. Natl. Acad. Sci. 114. 10.1073/pnas.1717421114.

8. Schadt, E.E., Monks, S.A., Drake, T.A., Lusis, A.J., Che, N., Colinayo, V., Ruff, T.G., Milligan, S.B., Lamb, J.R., Cavet, G., et al. (2003). Genetics of gene expression surveyed in maize, mouse and man. Nature 422, 297–302. 10.1038/nature01434.

9. Battle, A., Mostafavi, S., Zhu, X., Potash, J.B., Weissman, M.M., McCormick, C., Haudenschild, C.D., Beckman, K.B., Shi, J., Mei, R., et al. (2014). Characterizing the genetic basis of transcriptome diversity through RNA-sequencing of 922 individuals. Genome Res. 24, 14–24. 10.1101/gr.155192.113.

10. West, M.A.L., Kim, K., Kliebenstein, D.J., van Leeuwen, H., Michelmore, R.W., Doerge, R.W., and St. Clair, D.A. (2007). Global eQTL Mapping Reveals the Complex Genetic Architecture of Transcript-Level Variation in Arabidopsis. Genetics 175, 1441–1450. 10.1534/genetics.106.064972.

11. Zhang, X., Cal, A.J., and Borevitz, J.O. (2011). Genetic architecture of regulatory variation in *Arabidopsis thaliana*. Genome Res. 21, 725–733. 10.1101/gr.115337.110.

12. The GTEx Consortium (2020). The GTEx Consortium atlas of genetic regulatory effects across human tissues. Science 369, 1318–1330. 10.1126/science.aaz1776.

13. GTEx Consortium (2017). Genetic effects on gene expression across human tissues. Nature 550, 204–213. 10.1038/nature24277.

14. Kawakatsu, T., Huang, S.C., Jupe, F., Sasaki, E., Schmitz, R.J., Urich, M.A., Castanon, R., Nery, J.R., Barragan, C., He, Y., et al. (2016). Epigenomic Diversity in a Global Collection of Arabidopsis thaliana Accessions. Cell 166, 492–505. 10.1016/j.cell.2016.06.044.

15. Vu, V., Verster, A.J., Schertzberg, M., Chuluunbaatar, T., Spensley, M., Pajkic, D., Hart, G.T., Moffat, J., and Fraser, A.G. (2015). Natural Variation in Gene Expression Modulates the Severity of Mutant Phenotypes. Cell 162, 391–402. 10.1016/j.cell.2015.06.037.

16. Rockman, M.V., Skrovanek, S.S., and Kruglyak, L. (2010). Selection at linked sites shapes heritable phenotypic variation in C. elegans. Science 330, 372–376. 10.1126/science.1194208.

17. Zhou, Y., Zhang, Z., Bao, Z., Li, H., Lyu, Y., Zan, Y., Wu, Y., Cheng, L., Fang, Y., Wu, K., et al. (2022). Graph pangenome captures missing heritability and empowers tomato breeding. Nature 606, 527–534. 10.1038/s41586-022-04808-9.

18. The Genographic Consortium, Elhaik, E., Tatarinova, T., Chebotarev, D., Piras, I.S., Maria Calò, C., De Montis, A., Atzori, M., Marini, M., Tofanelli, S., et al. (2014). Geographic population structure analysis of worldwide human populations infers their biogeographical origins. Nat. Commun. 5, 3513. 10.1038/ncomms4513.

19. Yang, T., Liu, R., Luo, Y., Hu, S., Wang, D., Wang, C., Pandey, M.K., Ge, S., Xu, Q., Li, N., et al. (2022). Improved pea reference genome and pan-genome highlight genomic features and evolutionary characteristics. Nat. Genet 10.1038/s41588-022-01172-2.

20. Peter, J., De Chiara, M., Friedrich, A., Yue, J.-X., Pflieger, D., Bergström, A., Sigwalt, A., Barre, B., Freel, K., Llored, A., et al. (2018). Genome evolution across 1,011 Saccharomyces cerevisiae isolates. Nature 556, 339–344. 10.1038/s41586-018-0030-5.

21. Brem, R.B., Yvert, G., Clinton, R., and Kruglyak, L. (2002). Genetic dissection of transcriptional regulation in budding yeast. Science 296, 752–755. 10.1126/science.1069516.

22. Legras, J.-L., Galeote, V., Bigey, F., Camarasa, C., Marsit, S., Nidelet, T., Sanchez, I., Couloux, A., Guy, J., Franco-Duarte, R., et al. (2018). Adaptation of S. cerevisiae to Fermented Food Environments Reveals Remarkable Genome Plasticity and the Footprints of Domestication. Mol. Biol. Evol. 35, 1712–1727. 10.1093/molbev/msy066.

23. Hughes, T.R., Marton, M.J., Jones, A.R., Roberts, C.J., Stoughton, R., Armour, C.D., Bennett, H.A., Coffey, E., Dai, H., He, Y.D., et al. (2000). Functional Discovery via a Compendium of Expression Profiles. Cell 102, 109–126. 10.1016/S0092-8674(00)00015-5.

24. Gasch, A.P., Spellman, P.T., Kao, C.M., Carmel-Harel, O., Eisen, M.B., Storz, G., Botstein, D., and Brown, P.O. (2000). Genomic expression programs in the response of yeast cells to environmental changes. Mol. Biol. Cell 11, 4241–4257. 10.1091/mbc.11.12.4241.

25. Duan, S.-F., Shi, J.-Y., Yin, Q., Zhang, R.-P., Han, P.-J., Wang, Q.-M., and Bai, F.-Y. (2019). Reverse Evolution of a Classic Gene Network in Yeast Offers a Competitive Advantage. Curr. Biol. CB 29, 1126–1136.e5. 10.1016/j.cub.2019.02.038.

26. Boocock, J., Sadhu, M.J., Durvasula, A., Bloom, J.S., and Kruglyak, L. (2021). Ancient balancing selection maintains incompatible versions of the galactose pathway in yeast. Science 371, 415–419. 10.1126/science.aba0542.

27. Poirey, R., Despons, L., Leh, V., Lafuente, M.-J., Potier, S., Souciet, J.-L., and Jauniaux, J.-C. (2002). Functional analysis of the Saccharomyces cerevisiae DUP240 multigene family reveals membrane-associated proteins that are not essential for cell viability. Microbiol. Read. Engl. 148, 2111–2123. 10.1099/00221287-148-7-2111.

28. Celińska, E., and Nicaud, J.-M. (2019). Filamentous fungi-like secretory pathway strayed in a yeast system: peculiarities of Yarrowia lipolytica secretory pathway underlying its extraordinary performance. Appl. Microbiol. Biotechnol. 103, 39–52. 10.1007/s00253-018-9450-2.

29. Gallone, B., Steensels, J., Prahl, T., Soriaga, L., Saels, V., Herrera-Malaver, B., Merlevede, A., Roncoroni, M., Voordeckers, K., Miraglia, L., et al. (2016). Domestication and Divergence of Saccharomyces cerevisiae Beer Yeasts. Cell 166, 1397–1410.e16. 10.1016/j.cell.2016.08.020.

30. Shobayashi, M., Ukena, E., Fujii, T., and Iefuji, H. (2007). Genome-Wide Expression Profile of Sake Brewing Yeast under Shaking and Static Conditions. Biosci. Biotechnol. Biochem. 71, 323–335. 10.1271/bbb.60190.

31. Pérez-Ortín, J.E., Querol, A., Puig, S., and Barrio, E. (2002). Molecular characterization of a chromosomal rearrangement involved in the adaptive evolution of yeast strains. Genome Res. 12, 1533–1539. 10.1101/gr.436602.

32. García-Ríos, E., Nuévalos, M., Barrio, E., Puig, S., and Guillamón, J.M. (2019). A new chromosomal rearrangement improves the adaptation of wine yeasts to sulfite. Environ. Microbiol. 21, 1771–1781. 10.1111/1462-2920.14586.

33. Stuecker, T.N., Scholes, A.N., and Lewis, J.A. (2018). Linkage mapping of yeast cross protection connects gene expression variation to a higher-order organismal trait. PLOS Genet. 14, e1007335. 10.1371/journal.pgen.1007335.

34. Lee, T.J., Liu, Y.-C., Liu, W.-A., Lin, Y.-F., Lee, H.-H., Ke, H.-M., Huang, J.-P., Lu, M.-Y.J., Hsieh, C.-L., Chung, K.-F., et al. (2022). Extensive sampling of *Saccharomyces cerevisiae* in Taiwan reveals ecology and evolution of predomesticated lineages. Genome Res., genome;gr.276286.121v2. 10.1101/gr.276286.121.

35. Duan, S.-F., Han, P.-J., Wang, Q.-M., Liu, W.-Q., Shi, J.-Y., Li, K., Zhang, X.-L., and Bai, F.-Y. (2018). The origin and adaptive evolution of domesticated populations of yeast from Far East Asia. Nat. Commun. 9, 2690. 10.1038/s41467-018-05106-7.

36. Martin, M. (2011). Cutadapt removes adapter sequences from high-throughput sequencing reads. EMBnet.journal 17, 10. 10.14806/ej.17.1.200.

37. Trapnell, C., Pachter, L., and Salzberg, S.L. (2009). TopHat: discovering splice junctions with RNA-Seq. Bioinforma. Oxf. Engl. 25, 1105–1111. 10.1093/bioinformatics/btp120.

38. Li, H. (2011). Improving SNP discovery by base alignment quality. Bioinforma. Oxf. Engl. 27, 1157–1158. 10.1093/bioinformatics/btr076.

39. 39. Auwera, G. van der, and O’Connor, B.D. (2020). Genomics in the cloud: using Docker, GATK, and WDL in Terra First edition. (O’Reilly Media).

40. Dobin, A., Davis, C.A., Schlesinger, F., Drenkow, J., Zaleski, C., Jha, S., Batut, P., Chaisson, M., and Gingeras, T.R. (2013). STAR: ultrafast universal RNA-seq aligner. Bioinformatics 29, 15–21. 10.1093/bioinformatics/bts635.

41. Liao, Y., Smyth, G.K., and Shi, W. (2014). featureCounts: an efficient general purpose program for assigning sequence reads to genomic features. Bioinforma. Oxf. Engl. 30, 923–930. 10.1093/bioinformatics/btt656.

42. Paradis, E., and Schliep, K. (2019). ape 5.0: an environment for modern phylogenetics and evolutionary analyses in R. Bioinformatics 35, 526–528. 10.1093/bioinformatics/bty633.

43. Zheng, X., Levine, D., Shen, J., Gogarten, S.M., Laurie, C., and Weir, B.S. (2012). A high-performance computing toolset for relatedness and principal component analysis of SNP data. Bioinformatics 28, 3326–3328. 10.1093/bioinformatics/bts606.

44. Love, M.I., Huber, W., and Anders, S. (2014). Moderated estimation of fold change and dispersion for RNA-seq data with DESeq2. Genome Biol. 15, 550. 10.1186/s13059-014-0550-8.

45. Handcock, M.S., Hunter, D.R., Butts, C.T., Goodreau, S.M., and Morris, M. (2008). statnet: Software Tools for the Representation, Visualization, Analysis and Simulation of Network Data. J. Stat. Softw. 24, 1548–7660. 10.18637/jss.v024.i01.

46. Langfelder, P., and Horvath, S. (2008). WGCNA: an R package for weighted correlation network analysis. BMC Bioinformatics 9, 559. 10.1186/1471-2105-9-559.

47. Müllner, D. (2013). **fastcluster** : Fast Hierarchical, Agglomerative Clustering Routines for *R* and *Python*. J. Stat. Softw. 53. 10.18637/jss.v053.i09.

48. Langfelder, P., Zhang, B., and Horvath, S. (2008). Defining clusters from a hierarchical cluster tree: the Dynamic Tree Cut package for R. Bioinformatics 24, 719–720. 10.1093/bioinformatics/btm563.

49. 49. The Gene Ontology Consortium, Carbon, S., Douglass, E., Good, B.M., Unni, D.R., Harris, N.L., Mungall, C.J., Basu, S., Chisholm, R.L., Dodson, R.J., et al. (2021). The Gene Ontology resource: enriching a GOld mine. Nucleic Acids Res. 49, D325–D334. 10.1093/nar/gkaa1113.

50. Russo, P.S.T., Ferreira, G.R., Cardozo, L.E., Bürger, M.C., Arias-Carrasco, R., Maruyama, S.R., Hirata, T.D.C., Lima, D.S., Passos, F.M., Fukutani, K.F., et al. (2018). CEMiTool: a Bioconductor package for performing comprehensive modular co-expression analyses. BMC Bioinformatics 19, 56. 10.1186/s12859-018-2053-1.

51. Korotkevich, G., Sukhov, V., Budin, N., Shpak, B., Artyomov, M.N., and Sergushichev, A. (2016). Fast gene set enrichment analysis (Bioinformatics) 10.1101/060012.

52. Castel, S.E., Levy-Moonshine, A., Mohammadi, P., Banks, E., and Lappalainen, T. (2015). Tools and best practices for data processing in allelic expression analysis. Genome Biol. 16, 195. 10.1186/s13059-015-0762-6.

53. Pockrandt, C., Alzamel, M., Iliopoulos, C.S., and Reinert, K. (2020). GenMap: ultra-fast computation of genome mappability. Bioinformatics 36, 3687–3692. 10.1093/bioinformatics/btaa222.

54. Lippert, C., Listgarten, J., Liu, Y., Kadie, C.M., Davidson, R.I., and Heckerman, D. (2011). FaST linear mixed models for genome-wide association studies. Nat. Methods 8, 833–835. 10.1038/nmeth.1681.

