## Supplemental Material for "Pan-transcriptome reveals a large accessory genome contribution to gene expression variation in yeast"

for

**This document includes:**

Supplementary figures S1-9

Supplementary figure legends

Descriptions for supplementary tables S1-9

Online Datafile descriptions

Figure S1

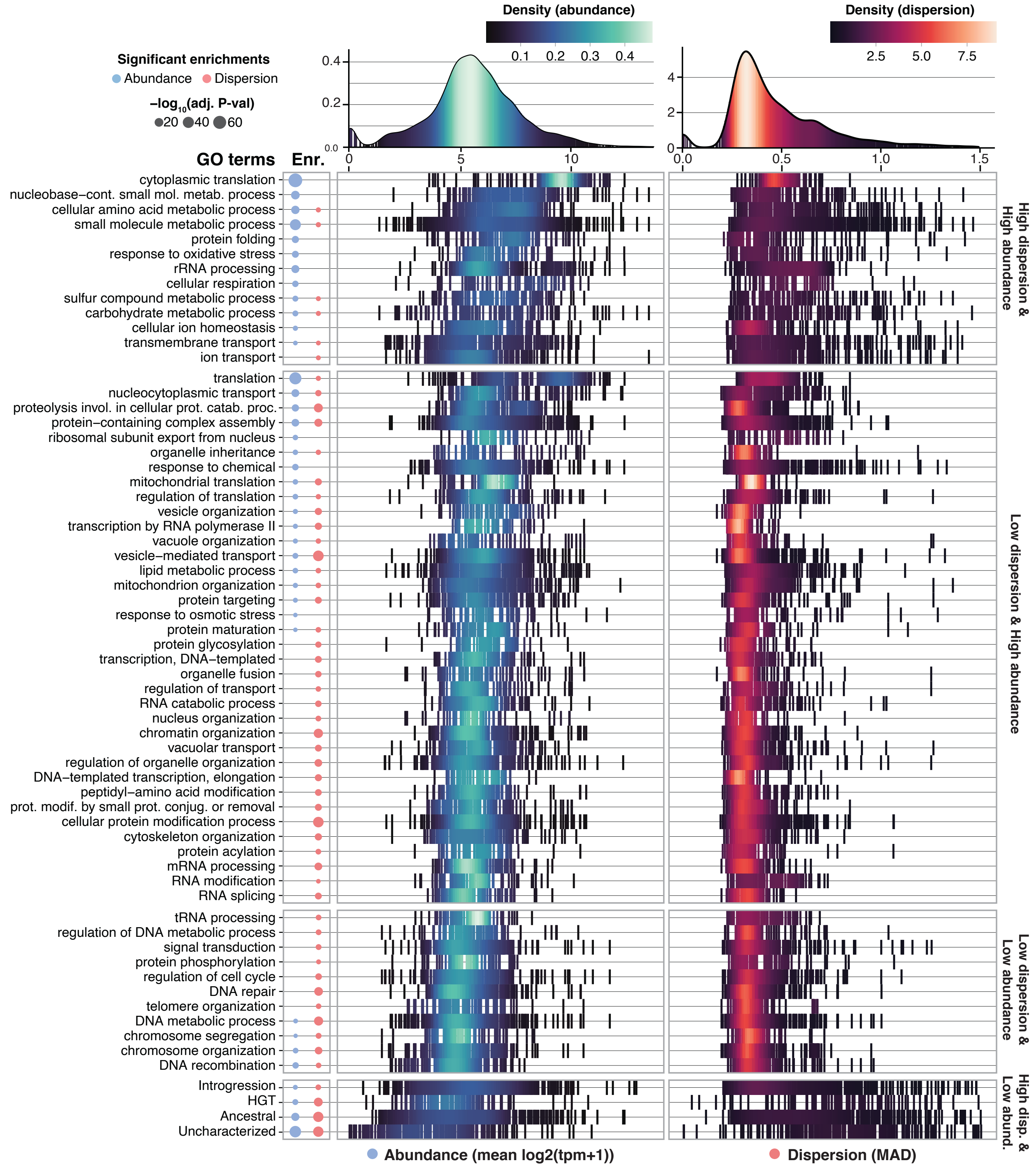


**Figure S1- Distributions and gene set enrichment analyses (GSEA) based on mean expression abundance and dispersion.** The global distribution for expression abundance, calculated as the mean log2 of TPM+1, and for expression dispersion, calculated as the mean absolute deviation, are shown on the top panel. Significant BP GO slim terms (59) and accessory gene subcategories (4) are indicated on the left. The strengths of the GSEA enrichment significance are indicated using the sizes of color-coded dots. The distributions of abundance and/or dispersion values for genes assigned in the corresponding terms are plotted. The distributions are grouped according to the quadrants as shown in Figure 2D. Within each group, terms are ranked based on the median expression abundance.

Figure S2

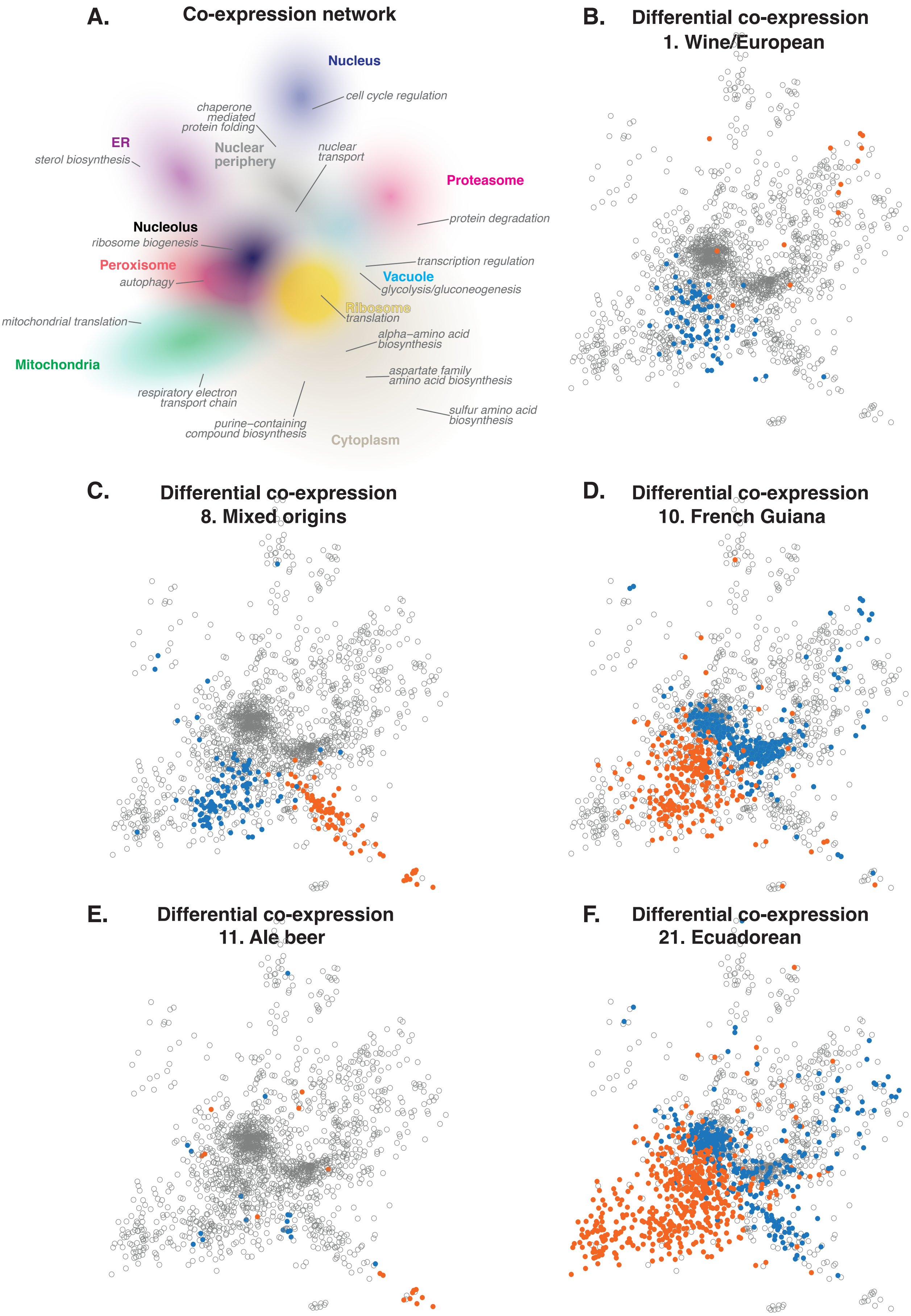

A.
Co-expression network
Nucleus
cell cycle regulation
chaperone mediated protein folding
ER
sterol biosynthesis
Nuclear periphery
nuclear transport
Proteasome
protein degradation
Nucleolus
ribosome biogenesis
Peroxisome
autophagy
transcription regulation
Vacuole
glycolysis/gluconeogenesis
Ribosome
translation
mitochondrial translation
Mitochondria
alpha–amino acid biosynthesis
aspartate family amino acid biosynthesis
respiratory electron transport chain
purine–containing compound biosynthesis
sulfur amino acid biosynthesis
Cytoplasm
B.
Differential co-expression
1. Wine/European
C.
Differential co-expression
8. Mixed origins
D.
Differential co-expression
10. French Guiana
E.
Differential co-expression
11. Ale beer
F.
Differential co-expression
21. Ecuadorean

**Figure S2- Differential co-expression by subpopulation. A.** Co-expression network annotations as shown in Figure 3A. **B-F.** Significant differential co-expressions in 1. Wine/European (**B**), 8. Mixed origins (**C**), 10. French Guiana (**D**), 11. Ale beer (**E**) and 21. Ecuadorean (**F**) subpopulations. Up-regulated genes are indicated in orange and down-regulated genes in blue. Modules and genes with significant co-expression changes by subpopulations are listed in Table S7 and recapitulated in Figure S3.

Figure S3

**A.**

| Subpopulations | #Isolates | Differential expression* #Down (in CO) | Differential expression* #Up (in CO) | Diff. co-expression† Down | Diff. co-expression† Up | Subpop-specific signatures Down | Subpop-specific signatures Up |
|---|---|---|---|---|---|---|---|
| 1. Wine/European | 354 | 310 (82) | 169 (14) | M4 | M5 | GO:0015758 | |
| 2. Alpechin | 16 | 143 (12) | 55 (3) | | | | |
| 3. Brazilian bioethanol | 33 | 182 (15) | 26 (5) | | | | GO:0019438 |
| 4. Mediterranean oak | 10 | 56 (1) | 8 (2) | | | | |
| 5. French dairy | 30 | 244 (23) | 188 (19) | | | GO:0016050 | GO:0006012 |
| 6. African beer | 17 | 175 (31) | 103 (5) | | | GO:0016050 | |
| 7. Mosaic beer | 18 | 18 (0) | 25 (2) | | | GO:0055085 | |
| 8. Mixed origin | 68 | 366(109) | 187 (79) | M4 | M7; M16 | GO:0006772 | |
| 9. Mexican agave | 7 | 49 (1) | 20 (1) | | | | |
| 10. French Guiana human | 30 | 696 (358) | 604 (266) | M1; M3; M5 | M2; M4; M6 | GO:0016050 | GO:0006772 |
| 11. Ale beer | 12 | 118 (17) | 96 (17) | | M16 | GO:0055085 | GO:0009313 |
| 12. West African cocoa | 13 | 35(3) | 17 (4) | | | | |
| 13. African palm wine | 26 | 155(4) | 35 (0) | | | GO:0046352 | |
| 14. CHNIII | 2 | 15 (0) | 4 (0) | | | | |
| 15. CHNII | 2 | 0 (0) | 0 (0) | | | | |
| 16. CHNI | 1 | n.a | n.a | | | | |
| 17. Taiwanese | 3 | 53 (5) | 17 (1) | | | GO:0006739 | |
| 18. Far East Asia | 9 | 67 (1) | 9 (0) | | | GO:0000723 | |
| 19. Malaysian | 5 | 97 (6) | 14 (0) | | | | |
| 20. CHN V | 2 | 23 (0) | 8 (0) | | | | |
| 21. Ecuadorean | 9 | 1007 (337) | 1015 (512) | M1; M3; M5; M7 | M2; M4; M6 | GO:0000723 | GO:0015758 |
| 22. Far East Russian | 4 | 226 (45) | 21 (4) | | | | |
| 23. North American oak | 13 | 156 (8) | 24 (0) | | | | |
| 24. Asian islands | 10 | 36 (3) | 19 (2) | | | | |
| 25. Sake | 46 | 243 (34) | 157 (8) | | | GO:0000723 | GO:0006772 |
| 26. Asian fermentation | 37 | 243 (54) | 105 (10) | | | | |
| M1. Mosaic region 1 | 17 | 0 (0) | 4 (0) | | | | |
| M2. Mosaic region 2 | 21 | 2 (0) | 0 (0) | | | | |
| M3. Mosaic region 3 | 110 | 18 (0) | 22 (0) | | | | |
| Unclustered | 43 | 59 (4) | 16 (0) | | | GO:0000723 | |

* Log2FoldChange > 0.3 & FDR < 0.05; † M1 ribosome biogenesis, M2 autophagy, M3 translation, M4 respiratory electron transport chain, M5 protein degradation, M6 mitochondrial translation, M7 alpha–amino acid biosynthesis, M16 sulfur amino acid biosynthesis.

**B.**

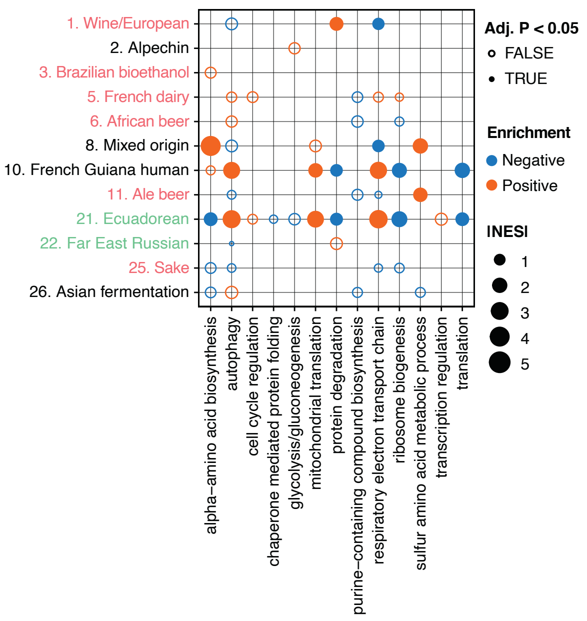


**C.**

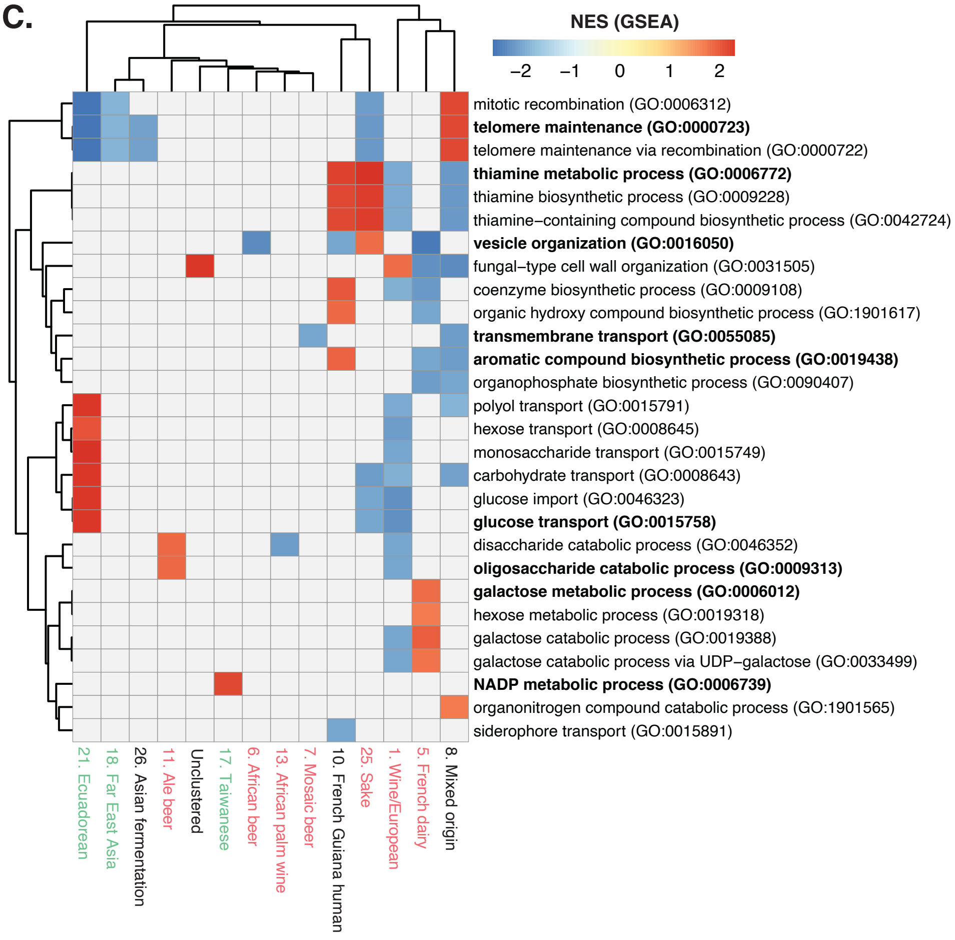

**Figure S3- Summary and functional enrichments of subpopulation specific differential (co-) expressions. A.** Overview of subpopulation size, number of significant differentially expressed genes and, enriched modules and biological processes (GO terms). Bars indicate the total number of differentially expressed genes, with darker shades representing the number of differentially expressed genes that overlap with the co-expression network. Full module names are indicated at the bottom of the plot. **B.** Module overrepresentation analyses (ORA) results by subpopulation. GSEA based normalized enrichment scores (NES) are shown. Dot sizes represent the absolute NES, with blue dots corresponding to underrepresentation (negative) and orange corresponding to overrepresented genes (positive). Solid dots indicate significant scores with adjusted P value $< 0.05$ (FDR) based 10,000 permutations. **C.** GSEA on log2 fold-change ranked differentially expressed genes by subpopulations. The heatmap is clustered based on the normalized enrichment scores across subpopulations and biological processes GO terms. Subpopulations with no enrichments are not shown. NES with an adjusted P-value $> 0.05$ (FDR) are masked in grey. FDR estimated based on 10,000 permutations.

Figure S4

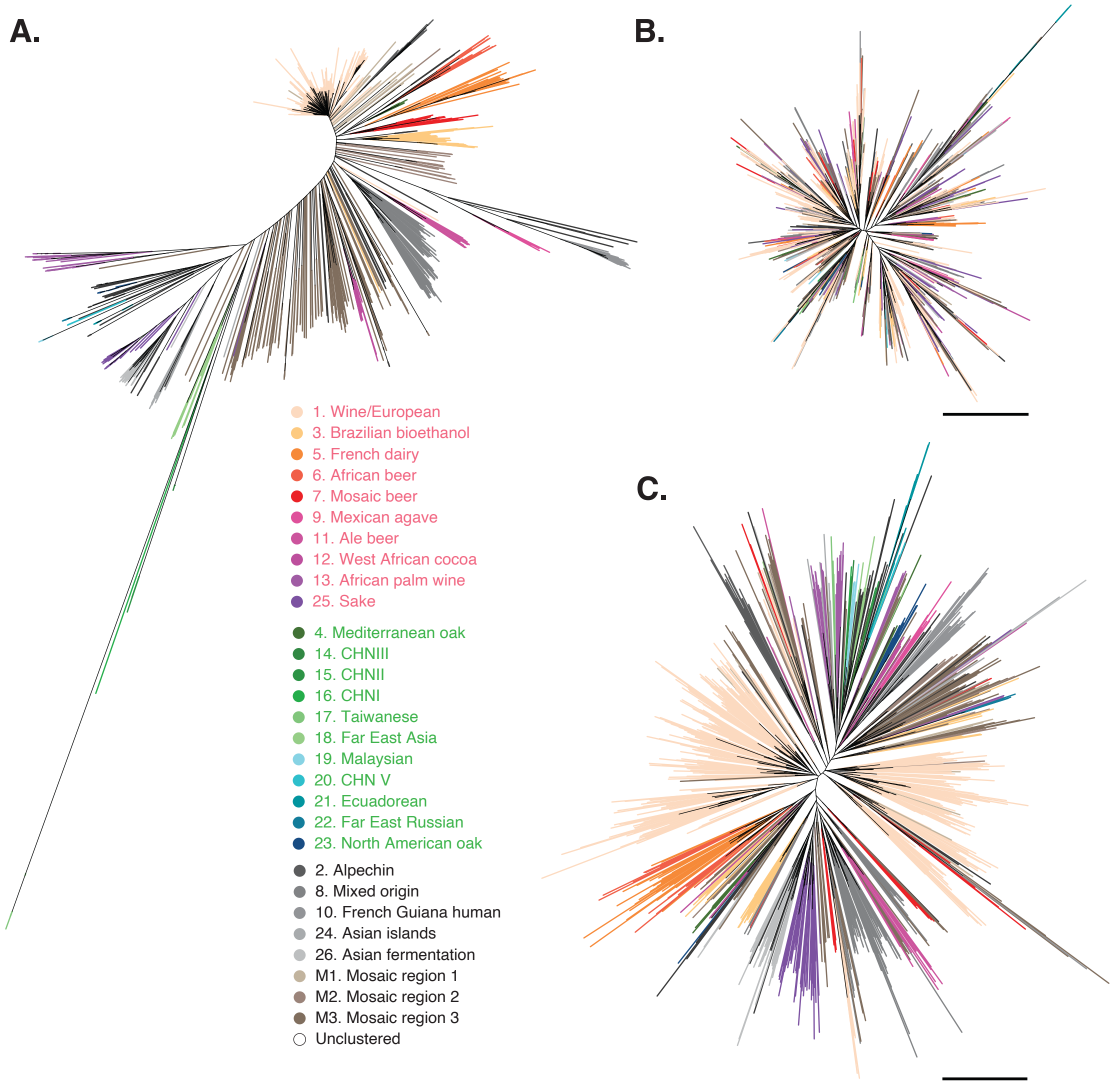


**Figure S4- Genetic divergence based on SNPs vs. gene expression diversity. A.** Neighbor-joining tree based on biallelic SNP positions across 969 isolates. Branches are color-coded according to the subpopulation definition as shown in Figure 1A. **B-C.** Neighbor-joining trees based on Euclidean distances using the 1,797 co-expression genes (**B**) and 2,209 differential expression genes (**C**). Scale bars are as indicated.

Figure S5

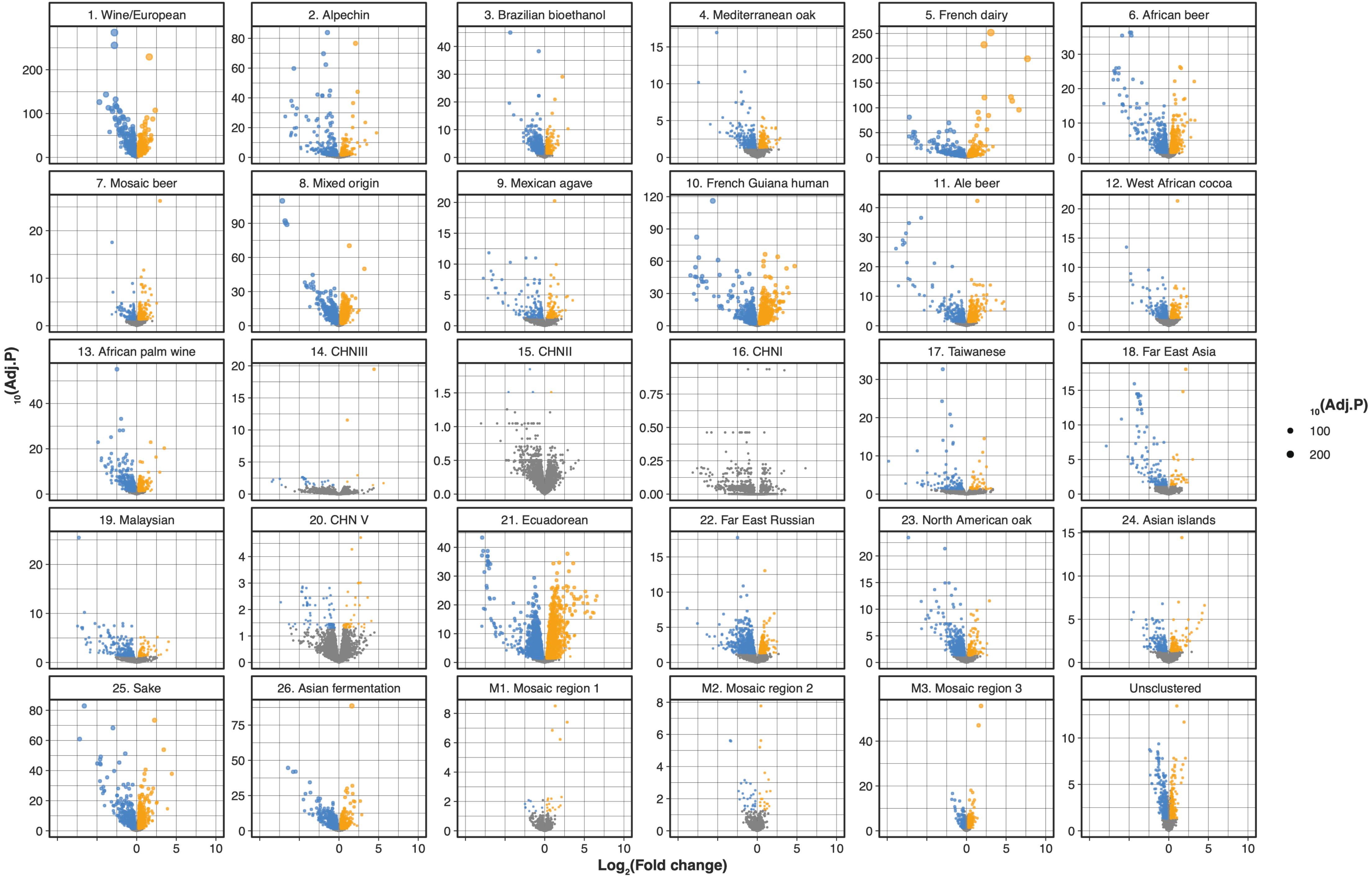
1. Wine/European
2. Alpechin
3. Brazilian bioethanol
4. Mediterranean oak
5. French dairy
6. African beer
7. Mosaic beer
8. Mixed origin
9. Mexican agave
10. French Guiana human
11. Ale beer
12. West African cocoa
13. African palm wine
14. CHNIII
15. CHNII
16. CHNI
17. Taiwanese
18. Far East Asia
19. Malaysian
20. CHN V
21. Ecuadorean
22. Far East Russian
23. North American oak
24. Asian islands
25. Sake
26. Asian fermentation
M1. Mosaic region 1
M2. Mosaic region 2
M3. Mosaic region 3
Unsclustered
10(Adj.P)
Log2(Fold change)
10(Adj.P)
100
200

**Figure S5- Volcano plots for differential gene expressions detected across each subpopulation.** Subpopulations are indicated on each subplot. For each subplot, x-axis corresponds to the log2 fold-change for each gene and y-axis correspond to the -log10 of the B-H adjusted P-values. Genes with an adjusted P-value < 0.05 and an absolute log2 foldchange > 0.3 are colored in blue for down-regulated genes and orange for up-regulated genes. Dot sizes are scaled to the y-axis.

Figure S6

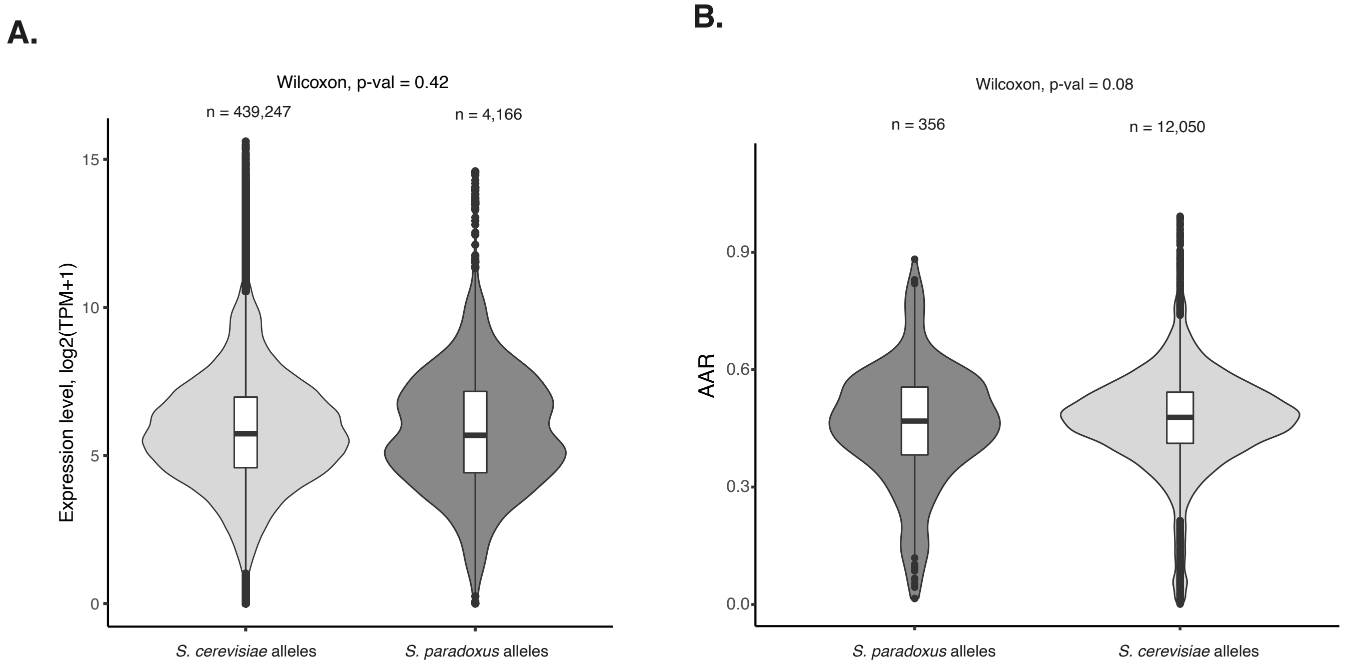


**Figure S6- Comparison of S. cerevisiae and S. paradoxus allele expression.**
**A.** Gene expression level comparison between 537 genes homozygous either for *S. cerevisiae* or *S. paradoxus* allele. The p-value is calculated using a two-sided Mann–Whitney–Wilcoxon test. **B.** Comparison between the alternative allele ratio (AAR) from heterozygous introgressed sites (after genetic allele balance filtration) and the AAR from non-introgressed heterozygous sites. The p-value is calculated using a two-sided Mann–Whitney–Wilcoxon test.

Figure S7

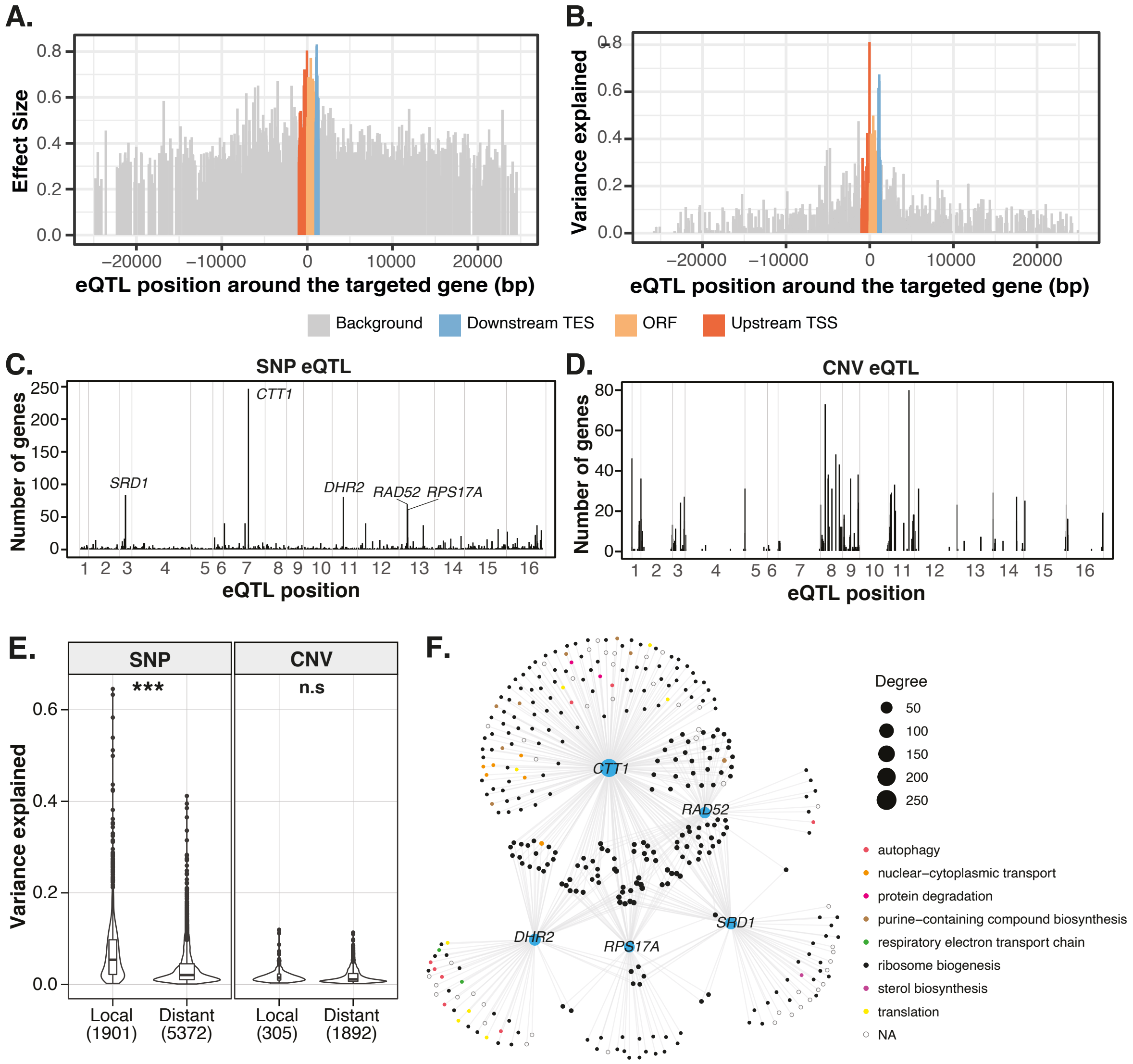


**Figure S7- eQTL feature summaries.**
**A-B.** Relative location of the local eQTL around the targeted gene and their effect sizes (**A**) and variance explained (**B**). The upstream region (red) corresponds to the 1,000 bp before the transcription start site (TSS), the downstream region (blue) corresponds to the 300 bp after the stop codon. The position of the eQTL located within the ORF (yellow) is scaled to 1,000 bp. The background regions contain the remaining regions located 25 kb before and after the gene. **C.** Hotspots for trans SNP-eQTL. The x-axis indicates the chromosomal locations for the eQTL and y-axis corresponds to the number of gene expression traits that are associated with a given eQTL. **D.** Hotspots for trans CNV-eQTL. The x-axis indicates the chromosomal locations for the eQTL and y-axis corresponds to the number of gene expression traits that are associated with a given eQTL. **E.** Comparison of the fraction of variance explained between local and distant eQTL for SNP and CNV eQTL types. Significant differences are indicated with stars. **F.** eQTL association network for top hotspots involving more than 50 expression traits. Size correspond to interaction degrees and expression genes are color-coded according to co-expression modules.

Figure S8

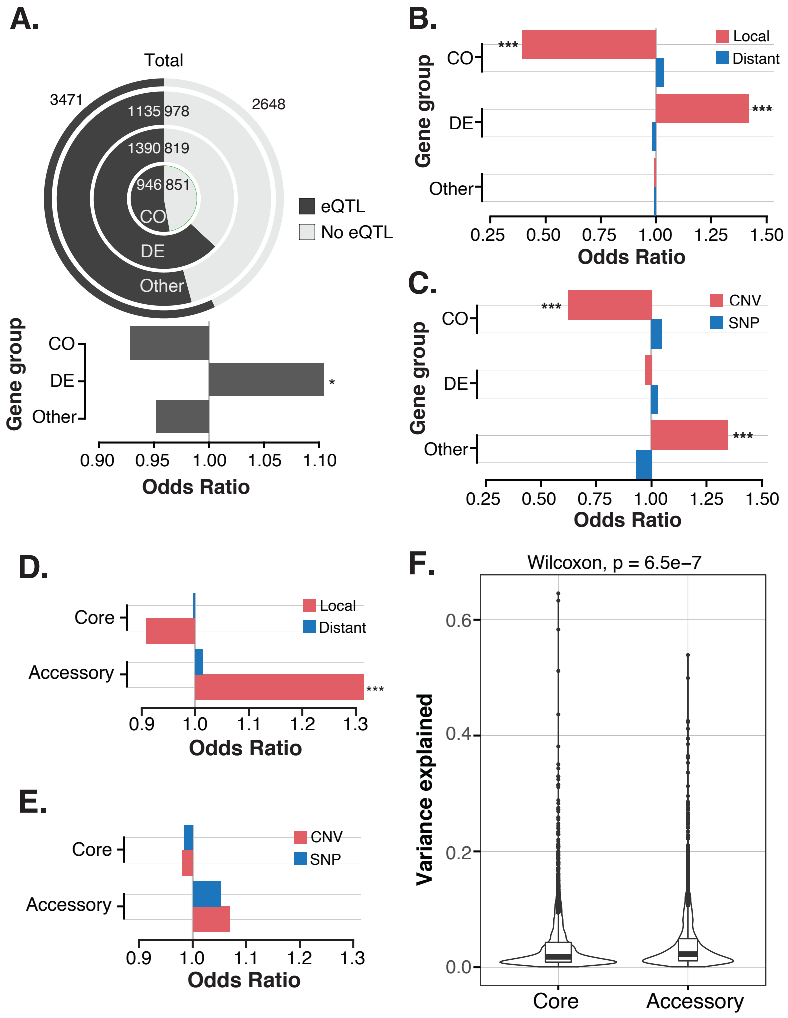


**FIgure S8- eQTL enrichments across different transcriptional network levels.**
**A.** From the inside out: the number and proportion of genes that are impacted by at least one eQTL for co-expression genes (CO), differential expression genes (DE), the remaining traits (Other) and the total dataset (outer ring). Fold-en- richments for CO, DE and Other genes on the proportion of genes impacted by at least one eQTL compared to the total set are presented as bars (odds ratio). Significant enrichment or depletion based on two-sided Fisher tests is indicated with stars. **B.** Fold-enrichments for CO, DE and Other genes on the proportion of genes impacted by at least one local (red) or distant (blue) eQTL com- pared to the total set. Odds ratios are indicated on the x-axis based on two-sided Fisher test. **C.** Fold-en- richments for CO, DE and Other genes on the proportion of genes impacted by at least one SNP (red) or CNV (blue) eQTL com- pared to the total set. Odds ratios are indicated on the x-axis based on two-sided Fisher test. **D.** Fold-enrichments for core and accessory genes on the proportion of genes impacted by at least one local (red) or distant (blue) eQTL compared to the total set. Odds ratios are indicated on the x-axis based on two-sided Fisher test. **E.** Fold-enrichments core and accessory genes on the proportion of genes impacted by at least one CNV (red) or SNP (blue) eQTL compared to the total set. Odds ratios are indicated on the x-axis based on two-sided Fisher test. **F.** Comparison of the fraction of variance explained between eQTL for accessory and core gene categories.

Figure S9

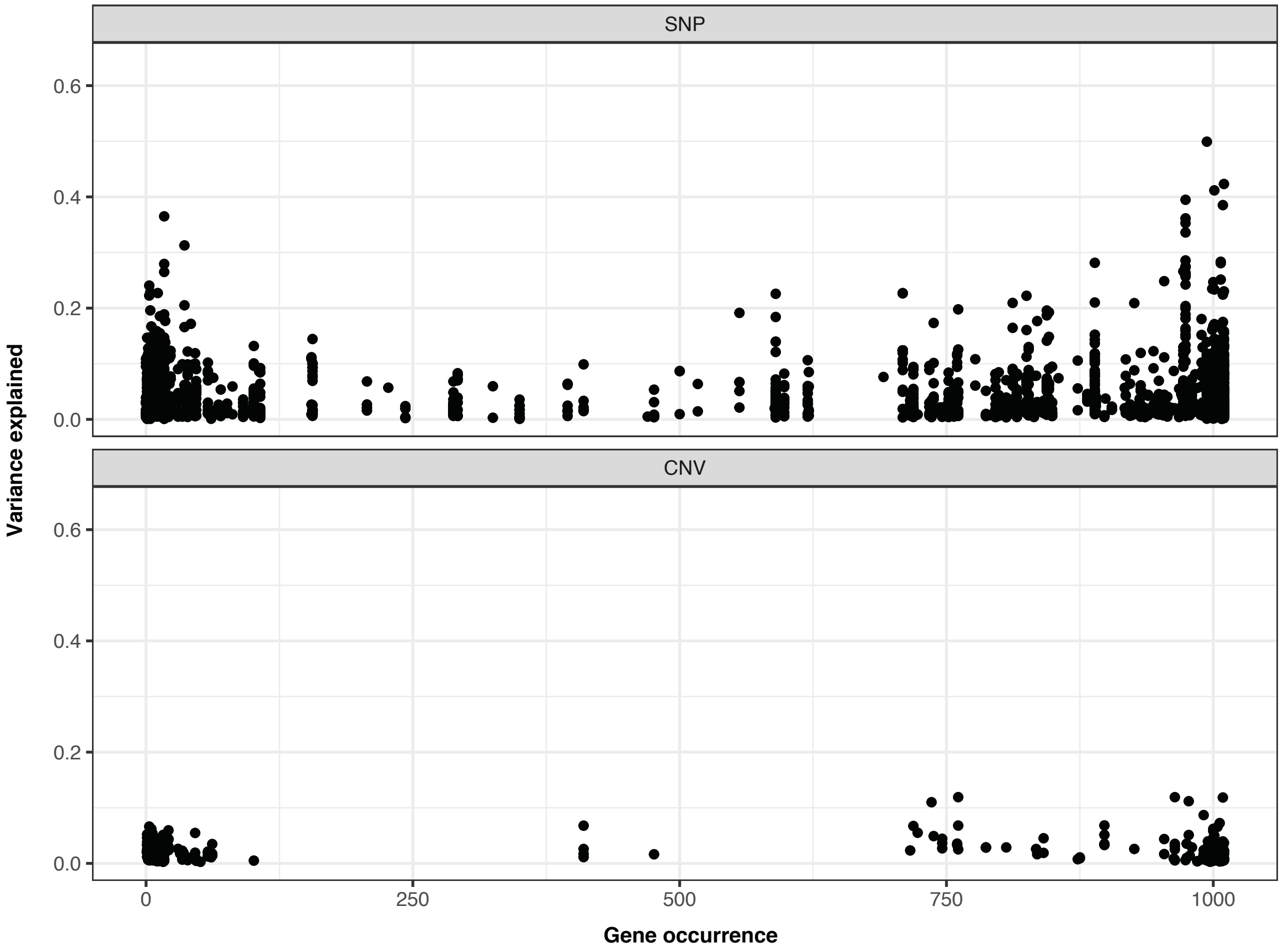


**Figure S9- Distribution of the fraction of variance explained per eQTL across accessory genes.** For each accessory gene associated with at least on eQTL, the fraction of variance explained (y-axis) is plotted against the number of isolates that carry the given accessory gene, i.e. gene occurrence, on the x-axis. The gene occurrence was obtained from the pangenome annotations across 1,011 isolates (Peter et al. 2018).

**Supplementary table descriptions**

**Table S1- Description of isolates included in this study.**

**Table S2- Description of genes included in this study.**

**Table S3- GSEA results on gene expression abundance and dispersion.**

**Table S4- Co-expression network module definition and connectivity**

**Table S5- GO term enrichment results across co-expression modules.** This table contains two tabs:

- BP: enrichments across GO biological processes;
- CC: enrichments across GO cellular compartments.

**Table S6- Significant differential gene expression across subpopulations.**

**Table S7- Differential co-expression by subpopulation based on module overrepresentation analyses**

**Table S8- GSEA results on differential gene expression by subpopulation**

**Table S9- eQTL enrichment analyses across transcriptional network levels.** This table contains three tabs:

- counts: summary counts for the types of eQTL associated with a given gene expression trait. Expression traits are grouped into co-expression (CO), differential expression (DE) and the remaining cases (Other).
- Fisher's test summary statistics for CO, DE and Other genes
- Fisher's test summary statistics for core and accessory genes

**Online Datafile descriptions**

**Datafile 1- Normalized gene expression levels (TPM) and raw counts data across 969 isolates**

**Datafile 2- GWAS results and statistics**
